# Distribution of mutation rates challenges evolutionary predictability

**DOI:** 10.1101/2022.12.12.518962

**Authors:** T. Anthony Sun, Peter A. Lind

**Affiliations:** Department of Molecular Biology, Umeå University, 90187 Umeå, Sweden; Umeå Centre for Microbial Research (UCMR), Umeå University, 90187 Umeå, Sweden

## Abstract

Natural selection is commonly assumed to act on extensive standing genetic variation. Yet, accumulating evidence highlights the role of mutational processes creating this genetic variation: to become evolutionarily successful, adaptive mutants must not only reach fixation, but also emerge in the first place, *i.e.* have a high enough mutation rate relative to the population size.

Here, we use numerical simulations to investigate how mutational biases impact our ability to observe rare mutational pathways in the laboratory and to predict outcomes in experimental evolution. We find the number of available mutations and the unevenness of their mutation rates to increase, sometimes impractically, the experimental effort required to discover all of them. Thus, most experimental studies lack power to directly observe the full range of adaptive mutations.

Modelling mutation rates as a distribution, we show that a substantially larger target size ensures that a pathway mutates more commonly. Therefore, we predict that commonly mutated pathways are conserved between closely related species, contrary to rare pathways. This approach formalizes our proposal that most mutations have a lower mutation rate than the average mutation rate measured experimentally. We suggest that the extent of genetic variation is overestimated when based on the average mutation rate.

## 1 Introduction

Natural selection acting on extensive standing genetic variation is usually considered the main determinant of adaptive evolution. This largely ignores biases in the origin of genetic variation, which may play a major role in guiding evolution (Yampolsky and Stoltzfus, 2001; Stoltzfus and McCandlish, 2017; Stoltzfus, 2021; Cano et al., 2022b,c; Monroe et al., 2022). Although it is debated in general (Svensson and Berger, 2019; Gomez et al., 2020; Svensson, 2022; Cano et al., 2022a), the importance of such biases is obvious when there are multiple genetic pathways to an adaptive phenotype with similar fitness. Practical consequences are likely on short time scales as well, for example in pathogen evolution, colonization of new ecological niches or in weak mutation, strong selection contexts in experimental evolution.

Biases in the production of genetic variation may result from two main causes. First, some genes may be more commonly observed to mutate simply because they have a larger number of mutations that can produce the phenotype, and thus represent a larger target size. For instance, when adaptive mutations may increase the expression of a gene either through a positive regulator or through a negative regulator, then evolution often promotes the latter since loss-of-function mutations are more common that gain-of-function mutations (Lind et al., 2015; Murray, 2020).

Secondly, differences in the rates of specific mutations leading to an adaptive phenotype may also produce a bias. Bacterial mutation rates vary from <10^-12^ gen^-1^ for specific deletions between distant endpoints without sequence homology (Koskiniemi et al., 2012), to 10^-3^ gen^-1^ for duplications between long sequence homologies (Anderson and Roth, 1981), with order-of-magnitude differences even for the same type of base pair substitution in the same genomic region (Sankar et al., 2016). Mutational hotspots are often the main determinant for the adaptive mutants found in experimental evolution studies, which suggests that they are relatively common (van Ditmarsch et al., 2013; Ferguson et al., 2013; Lind et al., 2016; Knöppel et al., 2018; Lind et al., 2019; Horton et al., 2021).

Experimental studies have measured mutation rates using two methods aiming at minimizing the effect of selection: mutation accumulation experiments, which are possible in a wide range of model organisms (Halligan and Keightley, 2009), and Luria-Delbrück fluctuation tests (Luria and Delbrück, 1943; Foster, 2006), designed specifically for microbes.

In mutation accumulation experiments, populations pass through small bottlenecks so as to reduce their effective size and let mutations fix largely independently from their fitness effect. The number and spectrum of mutations that have occurred over all generations can be determined using genome resequencing. Estimates of the average genomic mutation rate range from 0.8 × 10^-10^ to 5.3 × 10^-10^ bp^-1^ gen^-1^ for base pair substitutions, with the transition G:C>A:T being ten times more common than the transversion G:C>C:G, while insertion/deletion rates range from 0.2 × 10^-10^ to 2.0 × 10^-10^ bp^-1^ gen^-1^ (Lind and Andersson, 2008; Foster et al., 2015; Long et al., 2015; Sung et al., 2015; Dettman et al., 2016; Sung et al., 2016; Schroeder et al., 2017; Long et al., 2018; Pan et al., 2022). However, practical limitations imply that each of these estimates is typically based on an observation span of less than 100 mutations. This means that repeated mutations are not observed, and differences in the rate of specific mutations cannot be assessed.

In fluctuation tests, mutants are selected from independently grown microbial cultures. Estimations of the mutation rate to a particular phenotype, for example antibiotic resistance, relies on the Luria-Delbrück distribution for the number of mutants per culture (Luria and Delbrück, 1943; Foster, 2006). The spectrum of mutations can then be accessed by sequencing independent mutants and calculating the rate of each mutation as a fraction of the total mutation rate. Since the number of possible mutations to the adaptive phenotype is limited, it is possible to find repeated mutations and to measure differences in the rate of specific mutations. As mutation rates are not measured at the genomic level but often only in a single gene, the genomic context may impact the results. Fluctuation tests also tend to overestimate the average mutation rate per mutation since rare mutations are neither observed nor included in the number of possible mutations.

Here, we use numerical simulations to investigate how biases in the production of genetic variation affect evolution in the absence of differences in fitness between mutants. To account for the wide range of mutation rates, we model them as a distribution of mutation rates (DMR) rather than as a constant value. We only intend this theoretical model to shed light on the limitations of empirical studies. Combining insights from mutation accumulation experiments and fluctuation tests leads to the conclusion that a model DMR must have a higher average than median, because it has a long tail of rare high-rate mutations. This means that the mutation rate for most mutations is lower than traditionally assumed, as methods for estimating mutation rates are sensitive to extreme cases encountered in mutational hotspots.

We interpret those results using a well-established bacterial experimental evolution model system. Considering the evolution from a wild type to a phenotypically defined mutant, we show that characterizing the genotype-to-phenotype map (GPM) underlying this transition requires isolating a very large number of mutants when there are major differences in the number of possible mutations per gene. In particular, finding mutations in the same genes repeatedly is no reliable clue that other possible mutations are negligible, as long as the total number of mutants sequenced remains insufficient.

We also find that accurate experimental estimation of the genes’ relative mutational target size requires sequencing a much larger number of mutants than traditionally thought. Assuming a log-normal DMR, we show with a 95%-confidence interval that experimental data is compatible with a median mutation rate more than an order of magnitude lower than the average mutation rate. Finally, we show that the estimated log-normal DMR limits the predictability of experimental evolution on the level of individual mutations, but not at the level of genes, if there are many possible adaptive mutations per gene.

## 2 Model

### 2.1 Modelling approaches

We focus on the transition from the wild type to a mutant phenotype in a given species and consider mutation rates exclusively, as the origin of genetic diversity. In particular, we do not make any assumption regarding the interactions with selection impacting empirical data. Figure 1 provides an overview of our modelling approaches. We assume a fixed set of mutations and use two representations: mutation pie charts (MPCs) and a distribution of mutation rates (DMR).

**Figure 1:**
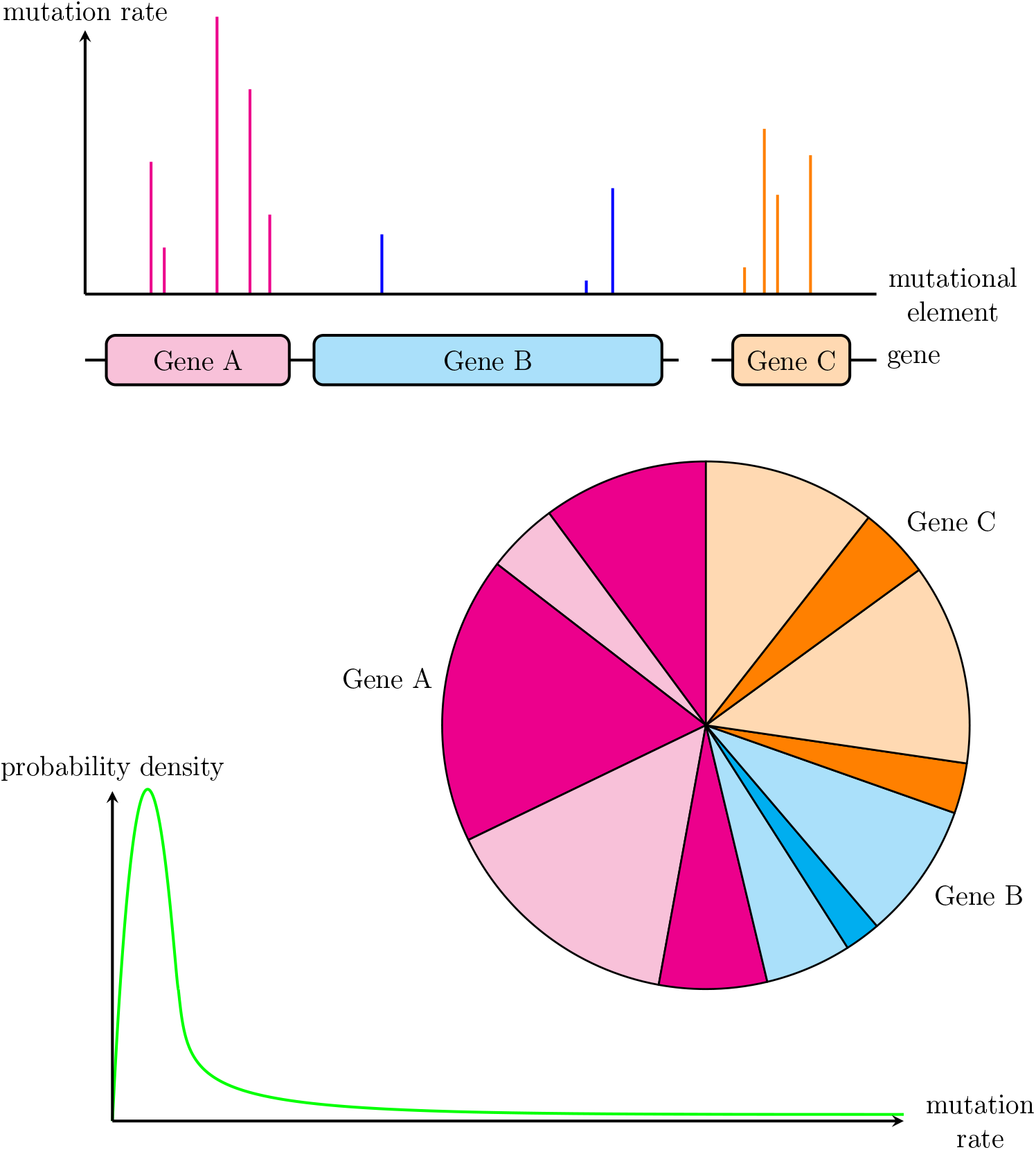
Approaches used. Graphical abstract of the approaches used in this article, showing the interpretation of the mutation rate for mutational elements defined on a molecular basis, the corresponding mutation pie chart (MPC) with relative mutation rates, as well as a conceptual distribution of mutation rates (DMR) considering absolute rates.

When focusing on the relative probability of each mutation, independently from absolute rates, it makes sense to use a pie chart representation: an MPC has probability 1, distributed among its shares. Each share accounts for one mutational unit, which may represent a mutation, a gene or a pathway. Thus, we model the observation of a mutation’s occurrence in the laboratory as a single, independent, random draw sampled from an MPC. In order to compare different MPCs, we introduce a metric summarizing it into a single number, the unevenness index *U*. It is commonly used in ecology and is based on Shannon’s diversity index. The unevenness index quantifies the global disbalance between the shares of the MPC, from perfect evenness, where all shares have the same size (*U* = 0), to maximal unevenness, where one share takes up the whole MPC (*U* = 1). Other metrics are possible (Appendix C.1).

When focusing on the absolute mutation rates however, we model the rate for each mutation as sampled from a DMR, which accounts for the variation in the rates of specific mutations. This allows us to investigate the effects of this variation on the repeatability of experimental evolution. It obviously makes sense to consider a DMR when an MPC has a large number of mutational elements. But interestingly, we can think of any MPC as a set of mutation rates sampled from a DMR. In particular, the same mutations sampling their rates twice independently from the same DMR to form two parallel MPCs can model closely related species.

We model the DMR as a log-normal probability distribution to satisfy several properties:

1. most mutation rates are low and very close to 0 (around 10^-10^ gen^-1^ for instance),
2. some are higher,
3. none is zero and almost none is extremely low (*e.g.* <10^-50^ gen^-1^).

Whether the support of the distribution covers all positive numbers or only the [0, 1] interval makes no difference in the context of mutation rates, where it suffices that values above 10^-2^ gen^-1^ are practically impossible. Among other distributions, such as the Gamma or the Beta distribution, the log-normal distribution is the only one to always satisfy condition 3., according to which any genetic sequence should in principle be able to mutate. The log-normal distribution is the distribution of the exponential of a normally distributed random variable. Its parameters *μ* and *σ* are the parameters of the underlying normal distribution.

### 2.2 Empirical data

To provide a direct link between theoretical results and laboratory experiments, we refer to the wrinkly spreader (WS) system in *Pseudomonas fluorescens* SBW25. One of the best characterized systems in experimental evolution, it has been used to address many fundamental questions in evolutionary biology (Rainey and Travisano, 1998; Kassen et al., 2000; Beaumont et al., 2009; McDonald et al., 2009; Hammerschmidt et al., 2014; Lind et al., 2015; Gallie et al., 2019; Lind et al., 2019) and can be extended to other *Pseudomonas* species to test evolutionary forecasts (Pentz and Lind, 2021). Here, we use an estimate of 500 mutations that can produce the WS phenotype, divided unevenly between 16 genes with 2 to 120 mutations each (Appendix B, Table 2). We also use experimental data from fluctuation tests for the *aws* operon within this system (Appendix B, Table 3).

When *P. fluorescens* is inoculated into static growth tubes, access to oxygen selects for mutants with an increased ability to colonize the air-liquid interface. The most successful type of such mutants is called *wrinkly spreader* because of its distinct colony morphology on agar plates. Single mutations characterize the GPM of the WS system, so that experimental data makes it possible to estimate the number of target genes and the number of mutations per gene: evidence reveals major differences, with prominent mutational hotspots biasing evolution towards certain genes (McDonald et al., 2009, 2011; Lind et al., 2015, 2019). Although WS mutants have a diverse genetic basis, they have similar fitness when competed against the ancestor (Lind et al., 2015) and can also be isolated independently from fitness effects (Lind et al., 2019). Thus, their evolutionary dynamics ideally fits our question since mutational phenomena play an important role compared to selection.

## 3 Results

### 3.1 Characterizing the genotype-to-phenotype map by completion experiments

Understanding the genotype-to-phenotype map (GPM) is fundamental to explain why evolution uses some routes rather than others. Experimentally however, it requires finding rare evolutionary pathways, sometimes orders of magnitude rarer than those commonly observed. As practical constraints limit how many mutants laboratory experiments can characterize, it is important to estimate the number of mutants required to find all genetic elements that can mutate to obtain a specific phenotype.

Repeatedly finding mutations in the same gene is often interpreted as an indication that the mutation pool has been exhausted. In order to explore theoretically to what extent this is valid, we draw independent, random observations from a known MPC, until all possible mutational elements have been drawn at least once. The result of such a completion experiment is the number of observations collected in order to achieve completion, simulating the number of mutants that a laboratory experiment would need to sequence.

We find that repeatedly finding the same mutations indeed indicates that the mutation pool has been exhausted, but only when the total number of observations is large enough. Simulations show that *large enough* can mean an impractically high number, especially when the possible mutations have very uneven mutation rates. Importantly however, the notion of *large enough* is not directly determined by the rarity of the rarest mutation.

#### 3.1.1 Reliability of experimental results depends on counts

The exact number of times a single type of mutation has been repeatedly observed, although it does not improve our molecular understanding of the phenotypic change, provides information on how reliably we picture available mutations. Indeed, Figure 2 confirms experimental studies suggesting that, by themselves, repeated observations do not necessarily mean that most genetic pathways have been discovered (Lind et al., 2015). For instance, the ideal case of the top panel shows a median at 52. This means that, at best, in half of the cases, three times as many observations as the actual number of pathways were necessary to get a complete picture of available mutational pathways. In some cases, an observation span of 200 sequenced mutants was still unable to reveal just 16 mutational elements: experimentalists would observe the same 15 mutations (or mutated genes) repeated over and over, on average >10 times each, without obtaining a single occurrence of the last one.

**Figure 2:**
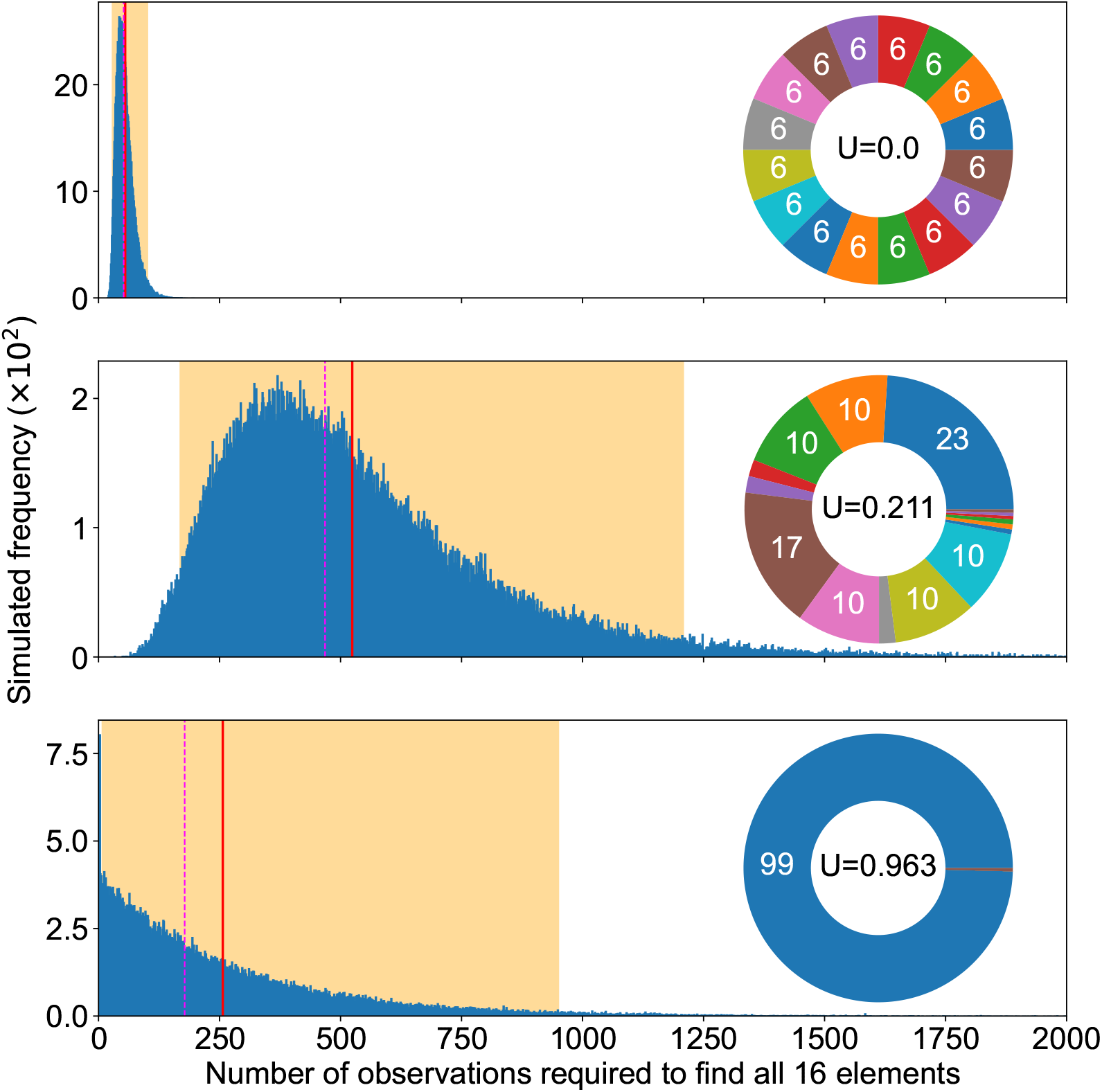
Completion experiments. Summary of completion experiments run on three examples of mutation pie charts (MPC) with different mutation rates in the even 16-case (top), the 16 pathways of the SW25 data (middle) and a case focusing on the rarest one in the SW25 data (bottom). The smallest number of replicates needed to find all mutational elements follows a skewed distribution (blue bars), with the highest simulated value being respectively 225, 3360 and 2906 out of 10^6^ experiments for each MPC. The average simulated value is shown as a solid red line (55, 524 and 257), the median as a dashed magenta line (52, 468, 178) and 95% of the experiment fell within the orange area ([29, 101], [169, 1208], [8, 950]).

This is because completion experiment results are distributed over a broad range. Thus, our picture of available mutational pathways heavily relies on luck, as long as we consider sequencing a limited number of mutants, which produces highly variable and thus unreliable results. As a consequence, it is essential to carefully record our observation span, *i.e.* the exact count of mutants experimentally sequenced, even if newly sequenced mutants do not reveal any new mutation.

#### 3.1.2 Rarest mutation alone does not capture completion speed

Counterintuitively perhaps, the number of experimental samples required to observe all possibilities does not critically rely on the rarest mutation pathway. Indeed, the middle and bottom panels of Figure 2 suggest that all mutational elements may influence our ability to complete the discovery of all possible mutations, not only the most uncommon ones. This suggests that the impact of the sheer number of mutational elements overrides matters of relative probability.

#### 3.1.3 Unevenness slows down completion

Beside the observation span, MPC-evenness makes it easier to achieve completion quickly, so that the even case provides a best-case scenario. Indeed, the two upper panels in Figure 2 show that the more uneven the MPC, the more observations we must collect to find all mutational elements. This applies to the expected number, to the median and to the variance. Whereas any other MPC increases the number of mutants to sequence as well as the results’ variability, the even case sets an ideal standard where completion experiments are the fastest and the most repeatably consistent.

In the best-case scenario of an even m-MPC, minimizing completion result with *m* available mutations (Appendix C.2.1), several formulas are available to inform the design of laboratory studies. The expected number of observations required to discover all mutational elements is approximately *m* (ln *m* + *γ* + 1/2m), where *γ* ≈ 0.577 is the Euler-Mascheroni constant. The variance is approximately *π*^2^*m*^2^/6 – (ln *m* + 1 + *γ*)*m* – 1/(12*m*). The expected number of additional observations required to observe a new, (*k*+1)-th mutation after discovering *k* different mutations is *m*/(*m* – *k*). And the maximum-likelihood estimation of the total number of mutations *m* given the number of elements already discovered *k* and the total sample size *obs* of observations gathered satisfies (Dawkins, 1991):

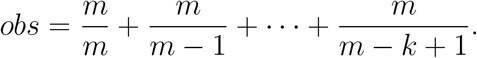

Let us illustrate how we can use this best-case scenario to generate information about a completion experiment. Suppose we are interested in a mutant phenotype produced by a number of mutations that we estimate to be *m* = 50. Were all mutation rates equal, so would we need to sequence 225 mutants on average to discover all 50 mutations. The standard deviation would be 62, meaning that, with some luck, an observation span of 300 mutants could suffice. Now, suppose we have already found 40 of the 50 mutations: we can expect to find a new one within 5 additional observations. All of that would be the best-case scenario: we must actually expect longer times in the likely case where mutation rates are uneven. But these simple formulas readily provide us the minimum observation span for an evolution experiment study to not be underpowered, *i.e.* allowing on average the observation of the mutation range.

### 3.2 Assessing the relative frequency of mutational elements

Related to the question of finding all genetic elements of the GPM, another important and experimentally more difficult problem is to quantify the probability that a certain genetic element mutates rather than another. This corresponds to estimating the MPC in terms of the relative share of each mutational element, which is a prerequisite for predicting evolution based on a model GPM.

As the number of observations increases to infinity, the law of large numbers almost surely ensures that we end up discovering all mutations, and that their relative frequencies converge to their relative mutation rates. Thus, we can theoretically access the shape of the MPC in an asymptotic manner. Does this mean that we need an infinite amount of experimental samples? No, luckily, things are more complex. Indeed, our simulations suggest that a few hundred observations are enough to converge to a correct estimation of a 16-MPC, on average and in median. Yet, luck plays an important role, so that certainty increases slowly with the observation span, especially if the MPC is very uneven.

#### 3.2.1 MPC-estimation converges fast in mean/median

Collecting more and more data improves our estimation from experimental samples at the level of the mean and at the level of the median. On the one hand, Figure 3 (top) shows that our attempts get closer and closer to a correct estimation of the real MPC as modelled by the unevenness: the expected error and the median error rapidly decrease when more mutations are observed, so that sequencing a few hundred mutants yields good approximations of the 16-MPC on average and in median.

**Figure 3:**
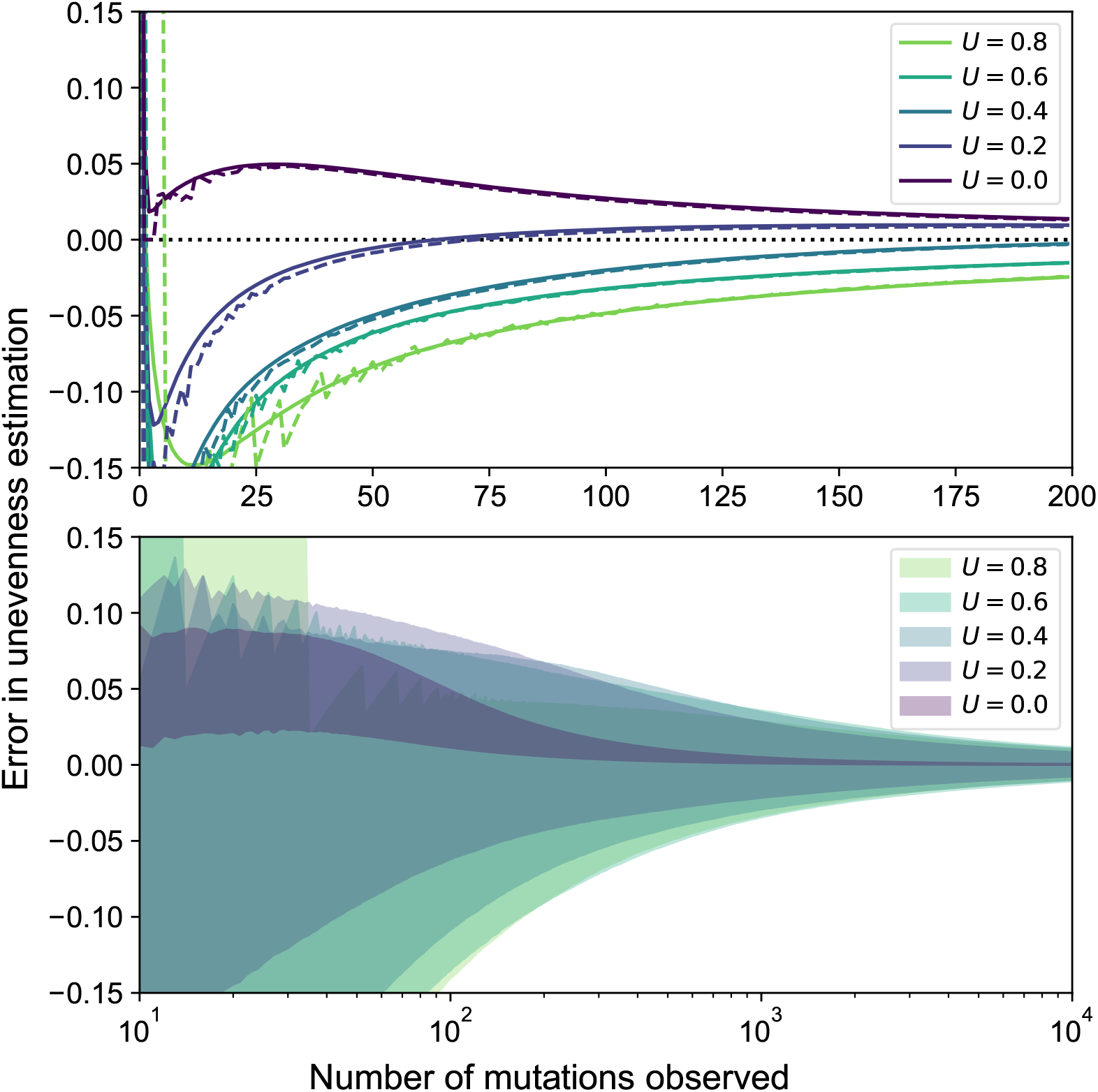
Convergence of MPC estimations. Impact of the real 16-MPC’s unevenness on the expected convergence of its estimations. Out of 10^5^ replicates for each unevenness value to estimate, central estimation errors are shown (top), with the average (solid) and median (dashed), as well as the 95% most common observed estimation errors (bottom).

#### 3.2.2 MPC-estimation converges slowly in variance

Collecting more data also improves our estimation from experimental samples when considering the variance of the estimation. Indeed, Figure 3 (bottom) suggests that the variability of the error shrinks. However, this variance drop happens on a much longer time scale: although the expected error and the median error are already low after sequencing 200 mutants from a 16-MPC, it remains very variable for a few more hundreds of mutants to collect. Importantly, after a high number of observations, the improvement of the expected error becomes negligible. For instance, after sequencing 1000 mutants from a 16-MPC, it would not be worth extending the observation span to 10 000 since the expected error would remain almost identical.

#### 3.2.3 Unevenness makes convergence more uncertain

Finally, Figure 3 suggests a moderate impact of the unevenness of the true MPC on our estimation when accumulating observations. The more even the true MPC, the faster the estimated MPC becomes correct. On the contrary, if the true MPC is more uneven, then we need to sequence more mutants in order to correctly estimate its unevenness. This does not hold for the trivial case where *U* = 1, since having a unique available mutation yields and immediate correct estimation of the MPC. These findings suggest that, in addition to finding all possible mutations, estimating their relative frequencies also requires an observation span of ideally several hundred mutants to sequence.

### 3.3 Viewing absolute mutation rates as a distribution

If mutation rates vary over orders of magnitude even for the same type of mutation (Sankar et al., 2016), then modelling them by a single average value may lead to erroneous conclusions. The concept of a distribution of mutation rates (DMR) can help think about mutation rate variations in a more nuanced way. So, we investigate how a DMR would affect our ability to predict evolutionary outcomes, exemplified by the WS model system.

#### 3.3.1 Log-normal DMR estimation from data

First, we calibrated a log-normal DMR using data from the WS system. Assuming that the mean was equal to the observed average of *E_obs_* = 5 × 10^-11^ gen^-1^ (Lind and Andersson, 2008; Foster et al., 2015; Long et al., 2015; Sung et al., 2015; Dettman et al., 2016; Sung et al., 2016; Schroeder et al., 2017; Long et al., 2018; Pan et al., 2022), we estimated the parameters of the log-normal DMR numerically based on the MPC-unevenness metric (Figure 4). We found that the data presented in Appendix B (Table 3) was compatible with values for parameter *σ* between 0.93 and 3.73 gen^-1^ (95%- CI). In the following, we use the maximum-likelihood estimate summarized in Table 1 as our calibrated mathematical tool, unless otherwise specified.

**Figure 4:**
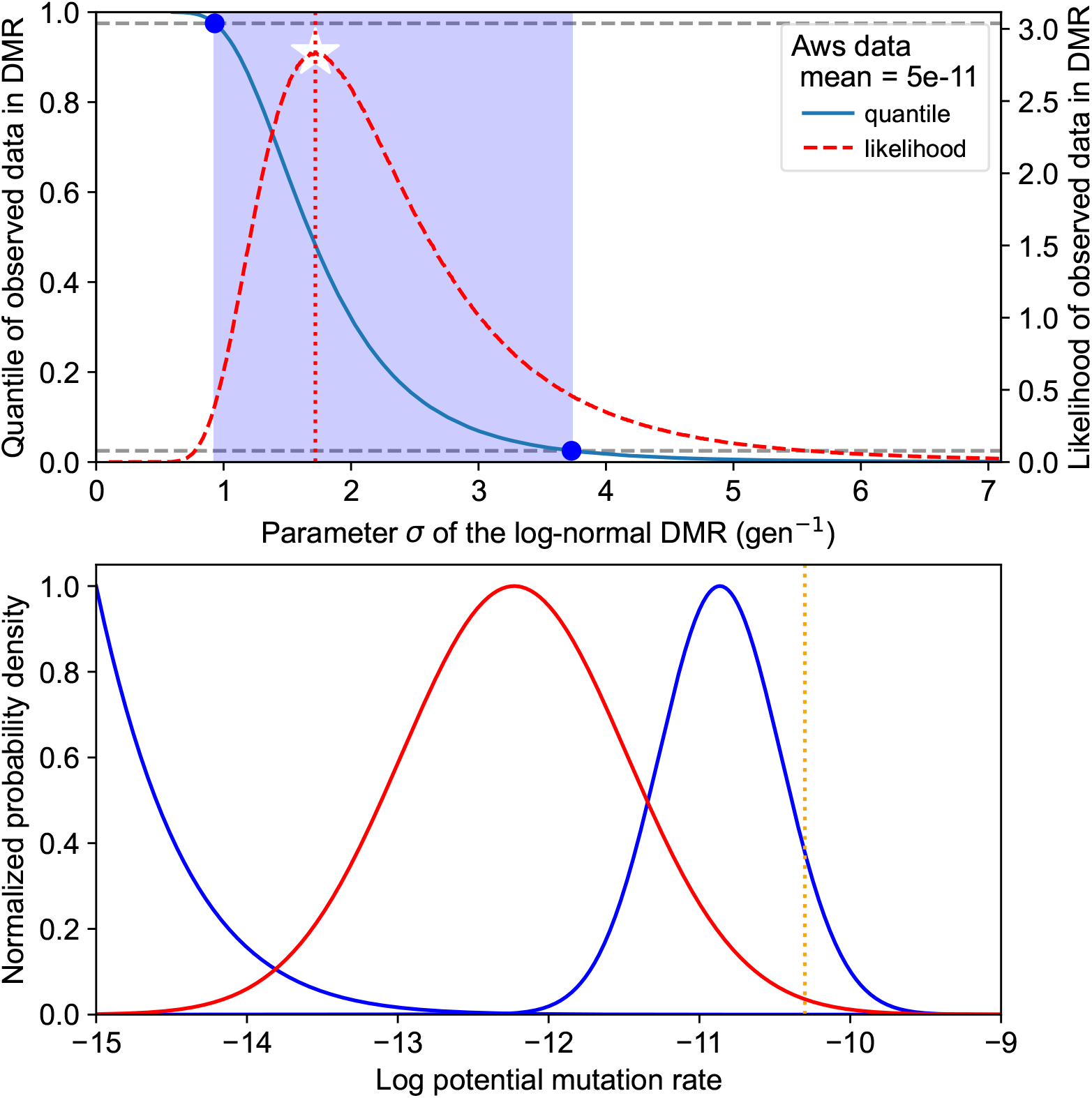
Log-normal DMR calibration approach. 95%-confidence interval (CI) for a log-normal DMR based on the experimentally informed assumption of average E_obs_ = 5 × 10^-11^ gen^-1^ and aws unevenness U_obs_ = 0.315. Top: Maximum likelihood (white star) for parameter σ and CI (blue band) constructed from the quantile curve crosses and the 95% range. 10^6^ 12-MPCs (aws setup) were sampled from the log-normal distribution with mean E_obs_ corresponding to each candidate value for σ. Bottom: Maximum likelihood estimation (red) for the DMR, lower (blue, left) and upper (blue, right) boundaries of the CI assuming mean E_obs_ (dashed orange line).

**Table 1:** Log-normal DMR calibration results. Unevenness-based maximum-likelihood estimation of the log-normal DMR from the data presented in Appendix B (Table 3).

| Characteristic | Value |
| --- | --- |
| parameter $\hat{\mu}$ | -25.20 |
| parameter $\hat{\sigma}$ | 1.72 |
| mean $E_{obs}$ | $5.00 \times 10^{-11} \text{ gen}^{-1}$ |
| median | $1.14 \times 10^{-11} \text{ gen}^{-1}$ |
| mode | $5.90 \times 10^{-13} \text{ gen}^{-1}$ |

Following empirical observations into assuming a skewed DMR implies that the average mutation rate does not correspond to the median, nor to the mode. Indeed, the data supports log-normal DMRs with a median ≤3.25 × 10^-11^ gen^-1^ and a mode ≤1.37 × 10^-11^ gen^-1^ (95%-CI), substantially lower than the assumed mean. Although they are sensitive to noise, medians should always be preferred when outliers are expected, since they are statistically stable. But the question is then how to interpret the average values obtained experimentally. Specifically, how does the DMR view of mutation rates inform the repeatability of experimental evolution experiments?

#### 3.3.2 Gene mutation rates repeatably reflect large discrepancies in target size

In the DMR framework, the extent to which experimental results may be repeatable, either within a single species or between specis with similar GPMs, depends on two opposing forces: the deterministic force of the MPC-setup representing mutational target size, which promotes repeatability; and the stochastic force of the heavy-tailed DMR representing the variability of mutation rates, which acts against repeatability. Here, we refer to mutations defined at the molecular level as *sites*, and to larger units at the gene or metabolic pathway level as *gene*, as *pathway* or as (*mutational) target*, so that the pathway rate is the sum of the rates for each of its mutation sites. This way, the number of mutation sites represents the size of a mutational target.

Our numerical simulations suggest that gene mutation rates reflect large discrepancies in target size. Indeed, even though the calibrated log-normal DMR allows for the mean to be 100-fold larger than the modal mutation rate, it was not able to overcome the determinism of common pathways having 25 times more sites than rare pathways (Figure 5). Thus, the MPC-setup determines the most common pathways in most cases, *i.e.* the fact that two mutational targets have substantially different sizes often ensures that the larger one has a significantly higher mutation rate. Here, *substantially* quantifies the extent to which repeatability of the most common pathways is robust to DMR-induced fluctuations.

**Figure 5:**
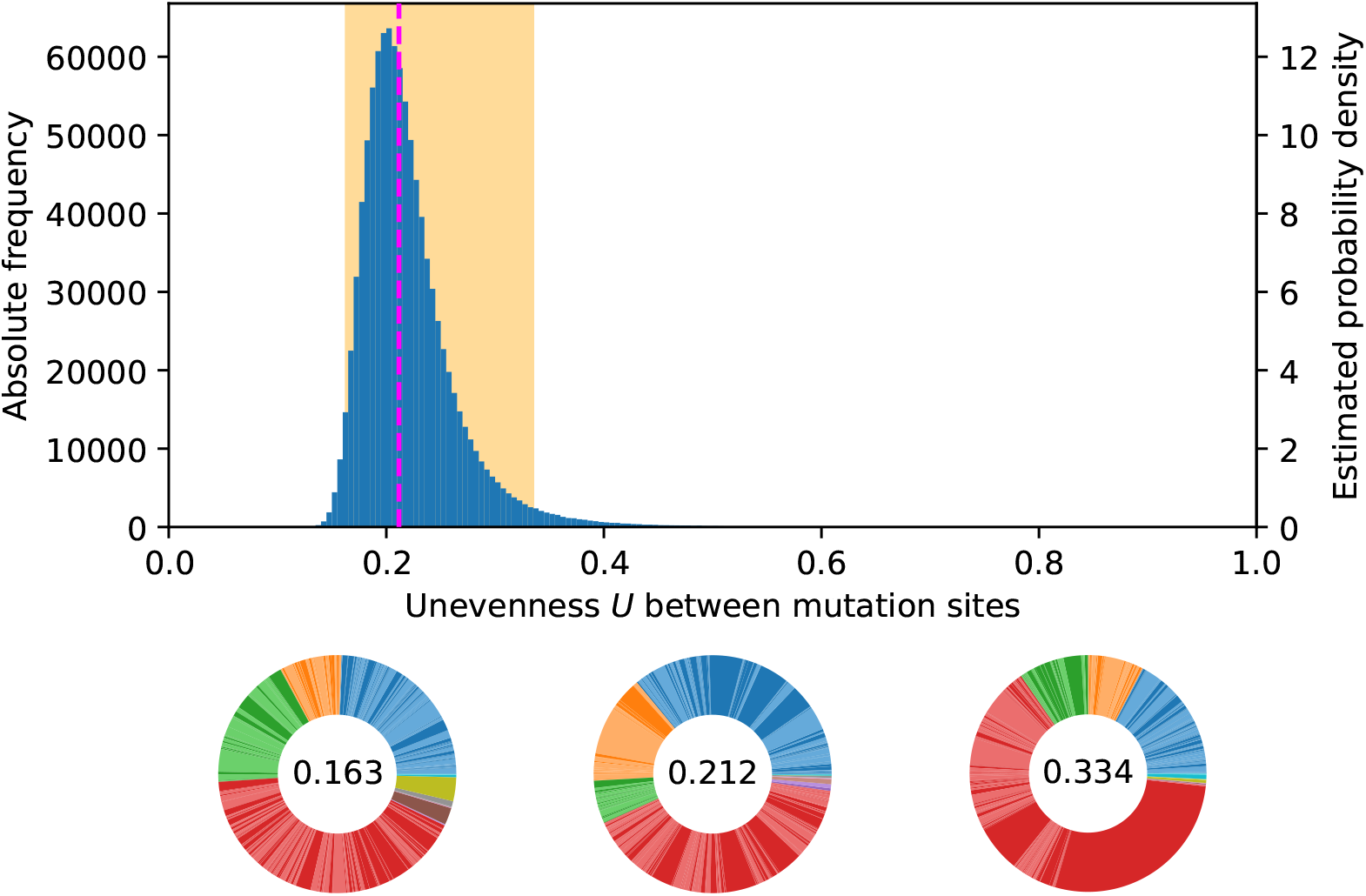
DMR’s impact on potential MPCs. Sampling 10^6^ MPCs of 500 mutations distributed among ten pathways as in the setup [145, 50, 50, 240, 2, 2, 2, 3, 3, 3] (corresponding to the data in Appendix B, Table 2) from a log-normal DMR with maximum likelihood estimated parameters μ = −25.15 and σ = 1.69 gives an unevenness distribution (blue) with median 0.212 (dashed magenta line) and 95% between 0.163 and 0.334 (shaded orange area). MPCs are shown for those three quantiles, each colour shade accounting for a mutation.

Which determinations of the MPC-setup are robust to the stochasticity resulting from the DMR? No analytical answer is known in the case of a log-normal DMR (Furman et al., 2020), but our numerical results suggest that a target size 15 times larger indeed ensures a higher effective mutation rate in practice (Figure 6). This result relies on realistic values for parameter *σ* of a log-normal DMR and 95 % certainty. Thus, the most heavy-tailed DMR compatible with the *aws* data does not blur the repeatability of the MPC-setup across orders of magnitude.

**Figure 6:**
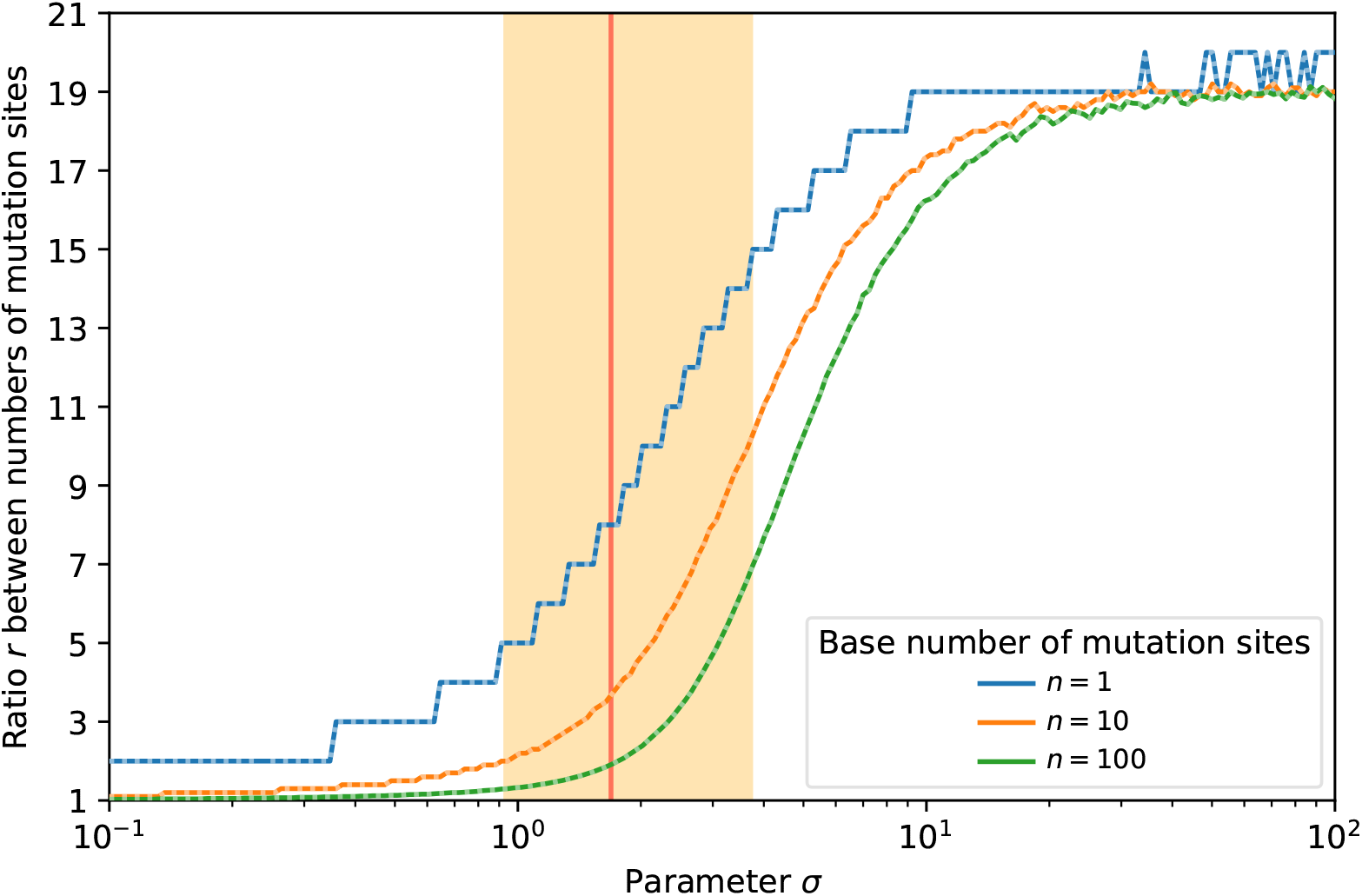
Robustness of MPC against DMR stochasticity. Consider two mutational pathways A and B of respective target sizes n and rn. Parameter σ of the log-normal DMR (i.e. its coefficient of variation) determines the threshold ratio r_thresh_ (line) under which >5% of cases challenge the robustness of the MPC-setup: in the region strictly below the lines, r is too low to ensure at 95% that A will have a higher mutation rate than B. The 5% average (solid coloured lines) and median (dotted white lines) cut-off was measured out of 10^6^ simulations in mean. The coloured area indicates the 95%-confidence interval for σ based on the aws data and the unevenness metric, with the maximum likelihood as a solid red line.

### 3.4 Exploring the impacts of the distribution of mutation rates

#### 3.4.1 Variability of MPC-unevenness from DMR models interspecific difference

We can interpret the variance of the MPC-unevenness distribution as a dimension of the potential difference between species. Then, the unevenness variability that a DMR produces models our ability to extend to closely related species the knowledge we have about one, assuming the same GPM. Assuming perfect knowledge of all mutation rates in one species, the narrowness of the unevenness distribution allows us to picture how much another species should be close in terms of the relative frequency of available mutations.

In Figure 5 for example, we mimic the data from Appendix B (Table 2) using the calibrated log-normal DMR from the *aws* data. Each mutation independently sampling its rate may result in more even (MPC on the left) or more uneven (MPC on the right) mutation rates. The median unevenness is 0.212, so that the data, with *U_obs_* = 0.315, suggests a relatively uneven observed MPC.

**Table 2:** Mutations to the wrinkly spreader phenotype. Estimates of the number of mutations for each gene from the smooth wild type to the wrinkly spreader phenotype in Pseudomonas fluorescens SBW25.

| Gene | Number of mutations |  |
| --- | --- | --- |
| *awsO* | 10 |  |
| *awsR* | 50 | total *aws*: 145 |
| *awsX* | 85 |  |
| *dgcH* | 50 |  |
| *mwsR* | 50 |  |
| *wspA* | 50 |  |
| *wspC* | 10 |  |
| *wspE* | 50 | total *wsp*: 240 |
| *wspF* | 120 |  |
| *wspR* | 10 |  |
| PFLU3448 | 2 |  |
| PFLU3571 | 2 |  |
| PFLU5960 | 2 |  |
| PFLU0956 promoter | 3 |  |
| PFLU1349 promoter | 3 |  |
| PFLU5698 promoter | 3 |  |
| Total | 500 |  |

#### 3.4.2 DMR variance increases MPC-unevenness, DMR mean has no impact

The variance of the DMR favours unevenness in resulting MPC, whereas its average has no impact (Figure 7). Indeed, the mean does not affect the MPC-unevenness distribution, since multiplying the rate of all mutations by the same factor does not impact their relative share in the MPC. However, the coefficient of variation of the DMR greatly impacts the unevenness: the higher the relative variance, the higher the unevenness. When the variance increases, the likely discrepancy between rare and frequent mutations becomes larger and the probability that few mutations dominate the MPC increases.

**Figure 7:**
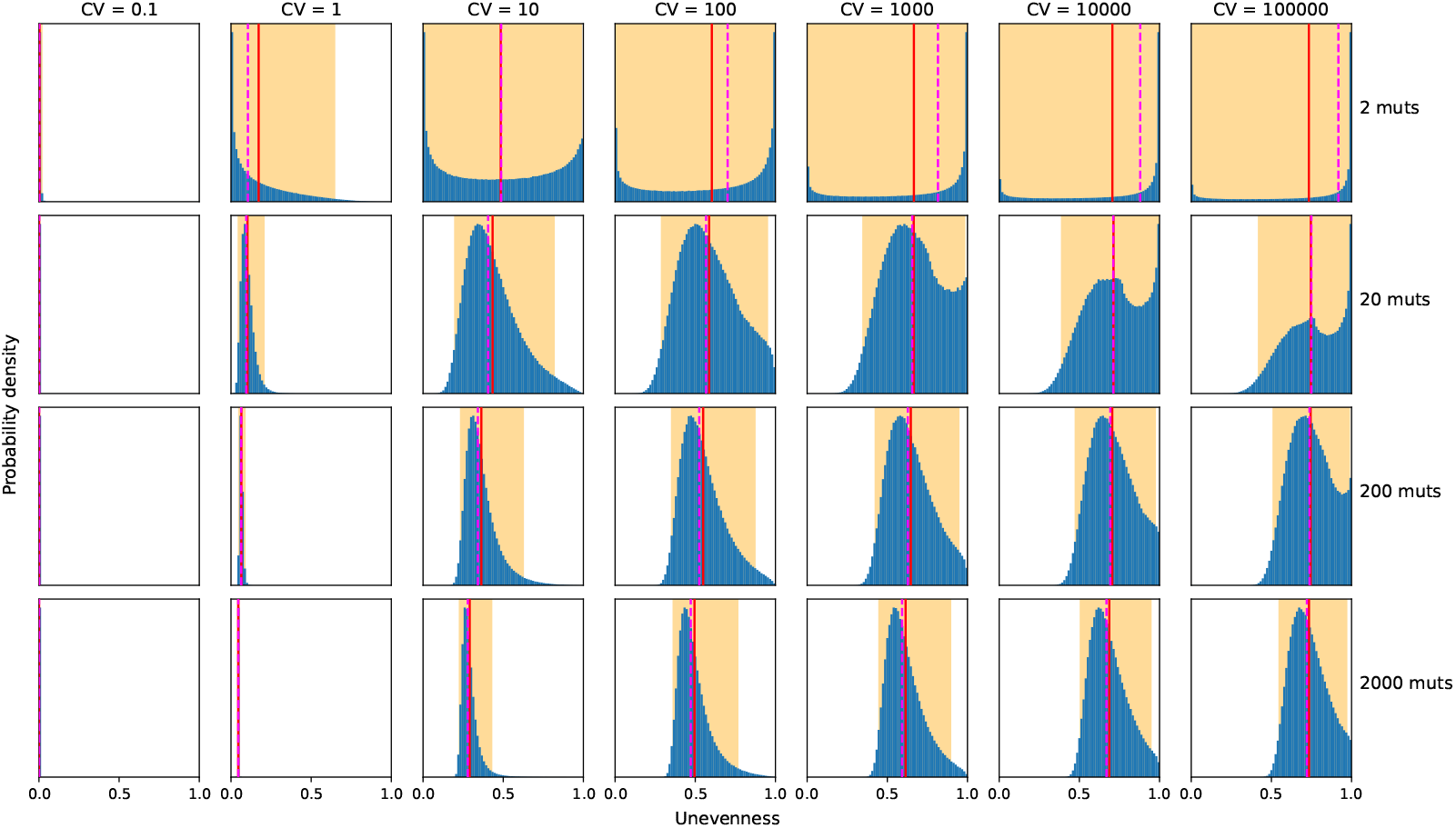
Impact of DMR on potential MPCs’ unevenness. Effect of the number of mutations and of the DMR’s coefficient of variation (CV) on the MPCs’ unevenness distribution. The examples shown use a log-normal DMR model to simulate 10^6^ replicated random MPCs in each panel.

#### 3.4.3 DMR shape impacts the absolute total mutation rate (TMR) when there are few mutation sites

The probability for a phenotype to appear in a population depends on the total mutation rate (TMR) from the wild type to that phenotype. The TMR is the sum of the rates for all mutations responsible for the mutant phenotype. Variation in the TMR between species could lead to major differences in experimental observations, for example the prediction that WS mutants should evolve or not in different species (Pentz and Lind, 2021). Assuming that the mutation rates independently follow a DMR with defined and finite expected value *r* and standard deviation *s,* the central limit theorem ensures that, as the number of mutations *n* increases, the distribution of the TMR converges to a normal distribution with location *nr* and variance ns^2^.

Therefore, whereas the TMR to phenotypes arising from a limited number of mutations largely depends on the exact shape of the DMR, an averaging phenomenon occurs in phenotypes resulting from a high number of mutations. This has an impact on the TMR distribution’s skewness (Figure 8 and Appendix C.3, Table 4). Indeed, since the DMR is skewed toward lower mutation rates, a TMR distribution resulting from few mutations is also skewed and has a median and a mode lower than the mean. On the contrary, having more mutations implies that the TMR tends to become symmetrically distributed with its median, mode and mean converging. This effect shows in the unevenness distribution since the unevenness of potential MPCs becomes less variable when more mutations are involved (Figure 7).

**Figure 8:**
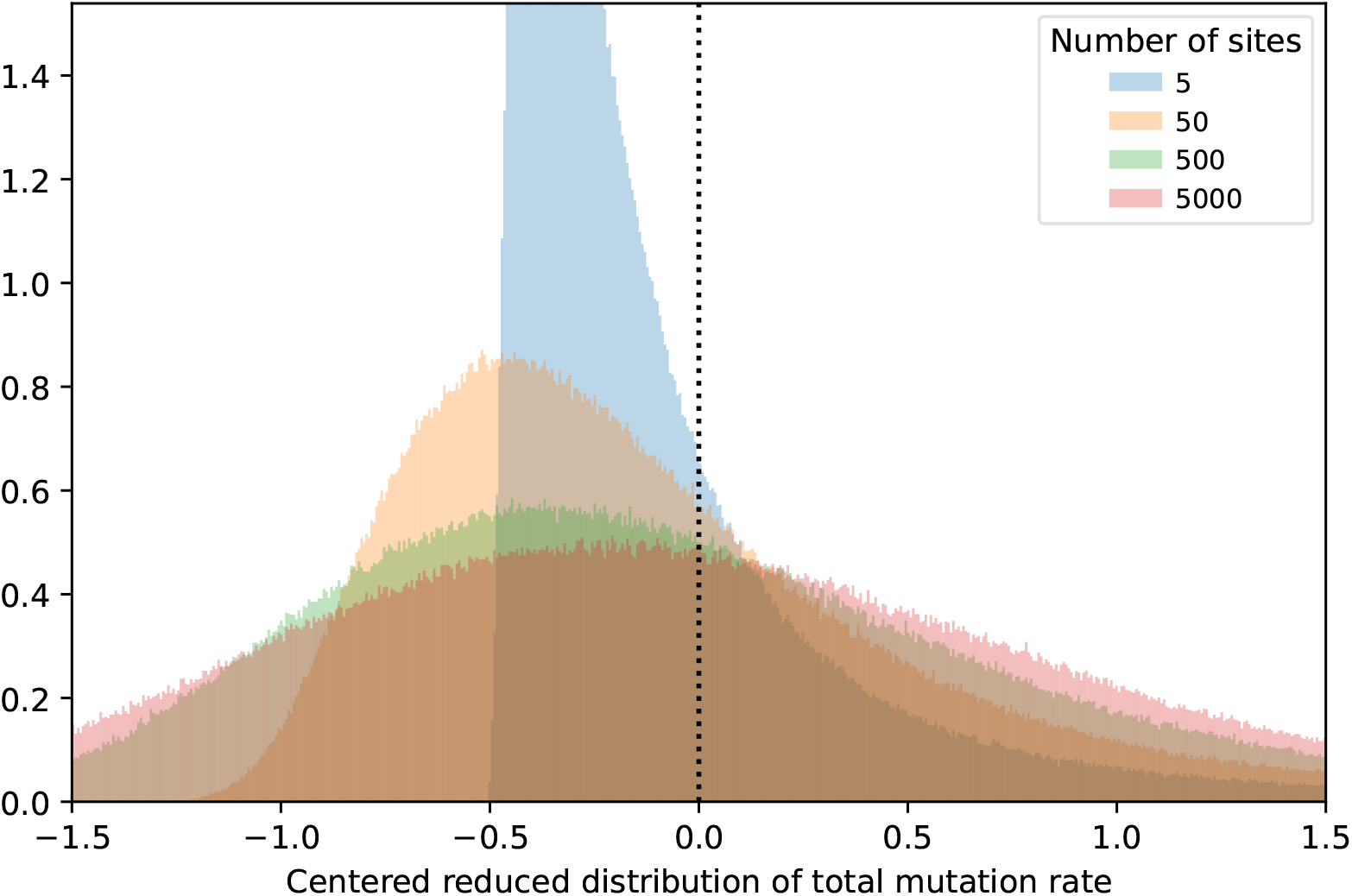
Distribution of the total mutation rate to the phenotype. Effect of the number of mutation sites on the total mutation rate, presented as a distribution centered around its mean and divided by its standard deviation (which would standardize a normal distribution). The examples shown use a log-normal DMR model to simulate 10^6^ for each number of mutation sites.

**Table 3:** Aws mutation spectrum. Occurence of aws mutations in a fluctuation test experiment on the transition from the smooth wild type to the wrinkly spreader phenotype in Pseudomonas fluorescens SBW25.

| Gene | Mutation | Occurrences | $U_{obs}$ |
| --- | --- | --- | --- |
| awsO | A 101 C | 1 |  |
|  | A 119 C | 1 |  |
| awsR | A 079 C | 9 |  |
|  | C 160 T | 1 |  |
|  | C 188 T | 1 |  |
|  | C 575 T | 2 |  |
|  | A 656 C | 1 |  |
| awsX | C 092 T | 1 |  |
| | Δ100-138 | 2 | |
|  | T 206 G | 1 |  |
| | Δ229-261 | 20 | |
|  | T 404 G | 1 |  |
| Total |  | 41 | 0.315 |

**Table 4:**
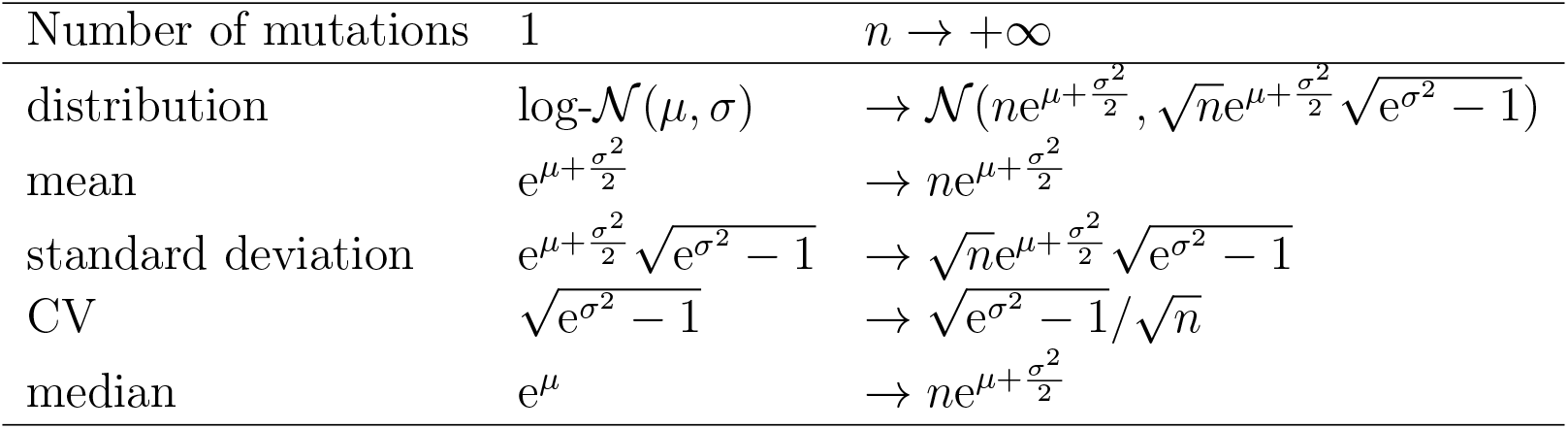
Impact of the number of mutations on distribution of the total resulting mutation rate (TMR).

### 4 Discussion

Although they constitute a fundamental evolutionary mechanism by introducing new genetic variation, mutations have traditionally been seen as simply providing the raw material for natural selection to act upon, with limited ability to influence the course of evolution (Stoltzfus, 2021; Cano et al., 2022b). However, biases in the origin of genetic variation may play a major role since they mean that frequencies at which particular mutants appear can differ by orders of magnitude, either in terms of phenotypes or in terms of genetic elements (Stoltzfus, 2021; Cano et al., 2022c).

#### 4.1 Relative mutation rates and how to deal with rare mutations

In this study, we used numerical simulations to explore how to take into account mutational biases and differences in the number of adaptive mutations per gene when interpreting empirical data, including results on evolutionary predictability. We showed that the number of possible adaptive routes are likely to be underestimated since most experimental studies, having an insufficient observation span, lack statistical power (Figure 2), which leads to an incomplete characterization of GPMs. Indeed, for the model system of the wrinkly spreader in *Pseudomonas*, directly finding all sixteen genes that can be mutated by a single mutation to cause a WS phenotype would require sequencing several hundred independent mutants, while accurately estimating their probability would require thousands of observations. Unevenness in the rates of available mutations scales up requirements beyond that of this theoretical, best-case scenario (Figure 2), rendering them impractical in laboratory settings.

Thus, the brute force approach of assuming that population sizes overcome the rarity of all mutations in microbiology experiments may be unable to reveal rare mutations in practice. A more strategical approach may be employed in cases where the most common pathways can be disabled (Lind et al., 2015), which has lead to the proposal of a hierarchy of mutational pathways in terms of negative regulators (commonly mutated), promoter mutations (intermediate) and rare activating mutations. Our results provide theoretical support for this view, because we showed that the speed at which we correctly picture the breakdown of mutation rates among pathways was short on average (Figure 3), in the range of laboratory experiments.

#### 4.2 Distribution of absolute mutation rates and repeatability of evolution

A large variation in mutation rates not only hinders the qualitative and quantitative discovery of mutational pathways, it is, additionally, a fundamental limitation for predicting evolutionary outcomes based on an understanding of GPMs (Lind et al., 2019). To illustrate this with an estimated GPM for the wrinkly spreader system in *Pseudomonas* (Pentz and Lind, 2021), we calibrated a log-normal DMR model using a small experimental dataset from *Pseudomonas fluorescens* SBW25 (Figure 4 and Table 1). It is unclear to what extent this data is representative for *Pseudomonas*, but the observed between-mutation ratio of 20 (Appendix B, Table 3) is similar to the ratio of 30 reported in *Pseudomonas aeruginosa* (MacLean et al., 2010). We also note the variation in mutation rates is likely to be higher than this experimental data suggests because possible mutations with lower rates are not observed experimentally, which leads to an overestimation of the average or median mutation rate per site.

We suggest that the DMR approach helps to disrupt thinking about mutation rates in terms of a single, constant value equated with the average (Figure 5). We found that it formalizes the fact that mean mutation rates are higher than median mutation rates (Figure 4 and Table 1). Therefore, most mutations have a lower rate than the average typically measured experimentally in mutation accumulation experiments. Put differently, natural selection acts upon a basis of genetic variation that is lower than traditionally assumed. Our calibrated log-normal DMR model allowed us to quantify that the median origin of genetic variation could actually be an order of magnitude below the mean mutation rate (Figure 4 and Table 1). This would extend to large populations of 10^9^ (Cano et al., 2022c) the range where mutational biases influence evolutionary outcomes, even though selection is the dominant factor in such cases.

Thus, evolutionary outcomes result from both a bias in variation origination (mutation) and a bias in variation fixation (selection). We should consider mutational biases as a key evolutionary driver beside selective processes because the standing variation selection acts upon is not as extensive as commonly assumed. A long history of origin-fixation models (McCandlish and Stoltzfus, 2014) supports this view, where evolutionary repeatability can be broken down into a contribution from selection and a contribution from mutation. The fixation step is often represented by random draws from a distribution of fitness effects. We cautiously suggest that a promising way forward may be to define the origin part of origin-fixation models as a DMR instead, which would allow for an estimation of the relative contribution of mutation versus selection. How this should be done exactly and how useful this approach may prove is an open matter for future studies.

Numerical results based on our calibrated model suggest that evolution may be predictable at the gene level in the following sense: pathways intrinsically more common by a factor 15 should indeed emerge more often in experimental studies. We can interpret this intrinsic predominance in terms of target size (Figure 6), but also in light of a molecular understanding (Lind et al., 2015). This provides theoretical and quantitative support to experimental studies exploiting the rate hierarchy of mutational pathways (Lind et al., 2015) in a forecasting perspective (Pentz and Lind, 2021).

Additionally, if we interpret a DMR as a null-model for repeatability within and between species repeatability, its variance (Figure 7) directly relates to the probability of parallel evolution (Stoltzfus, 2021), which we discuss further in Appendix C.4. Our findings suggest low levels of exact mutational parallelism between different species: while molecular evolution in a single species may be highly repeatable because of mutational hotspots, rare adaptive pathways with few possible mutations are likely to differ between species (Figure 5). Therefore, we predict that, when mutating to a wrinkly spreader phenotype, other *Pseudomonas* species will also use the main known pathways: *wsp, aws, mwsR* and *dgcH.* However, since each comprises many possible mutations, we can expect the particular mutations used for each to differ between species. We predict that genes with few possible mutations only appear common in a particular species where they happen to have a high mutation rate: we expect them to differ across species with little to no predictability. Moreover, since the wrinkly spreader phenotype is due to a large number of possible mutations, we can predict that the total mutation rate approximately follows a normal distribution across different species (Figure 8).

#### 4.3 Conclusions

We have argued that the mathematical shape of a DMR was a matter of modelling on the ground of first principles. Nevertheless, future experimental data on mutation rates could help improve the DMR model. Thus, we see a great need for more extensive experimental studies collecting and characterizing many independent mutants (Cano et al., 2022b). It would be especially interesting to study natural genomic contexts where multiple genes with large mutational target size present adaptive mutations.

Finally, we showed that information on MPCs, *i.e.* on relative mutation rates for the different mutations available, provides direct constraints on compatible DMRs. Therefore, finding ways to adequately measure and report the wide range of mutation rates, both in absolute and in relative terms, would generate empirical material to inform our model. Whether the mutation rate variation spectrum is similar across species remains to be discovered, as well as the ways in which the environment and DNA repair systems influence it. We hope to see much progress in the field since addressing these questions do not entail intrinsic technical difficulties. Thus, we encourage researchers to make experimental data available in an accessible format, so that the methods we developed here to calibrate a DMR model for mutation bias, based on such data, can easily transposed to other experimental models.

## 5 Methods and Materials

We ran numerical experiments in Python. Appendix A presents the rationale of their experimental design. Code is available at https://github.com/anthony-sun/Mutationrates.git.

We only use experimental data published previously, described in Appendix B.

## 6 Author contributions

TAS: Conceptualization, Methodology, Software, Validation, Formal analysis, Investigation, Data curation, Writing – original draft preparation, Writing – review & editing, Visualization.

PAL: Conceptualization, Methodology, Writing – original draft preparation, Writing – review & editing, Supervision, Funding acquisition.

## 7 Funding

This work was supported by grants from the Swedish Research Council (Vetenskapsrådet 2019-04859), Carl Trygger’s Foundation for Scientific Research (CTS 19:204) and the Åke Wiberg foundation (M18-0142) to PAL.

## 8 Acknowledgements

We thank Yashraj Chavhan for thoughtful comments on the manuscript.

## 9 Competing interests

The authors declare no competing interests.

## A Numerical appendix

### A.1 Completion experiments from known MPC

If we model the observation of each occurrence of a mutation in the laboratory as a single, independent, random draw sampled from an MPC, we can use the coupon collector’s problem to obtain theoretical results. Indeed, the non-equiprobable extension of this mathematical problem then tells us about the number of observations needed to discover all possible mutational pathways: its expected value is known exactly (Nath, 1974; Flajolet et al., 1992; Ferrante and Saltalamacchia, 2014) as well as its variance (Nath, 1974; Boneh and Hofri, 1989).

Yet, the exact mathematical formulas involved are too complicated to be useful in an experimental context. Indeed, laboratory studies need, on the one hand, working approximations rather than exact mathematical formulas for the variance and, on the other hand, an estimation of the median number of mutants to be collected rather than the expected number.

We propose a simple way to visualize the inherent variation that laboratory observations impose. Starting from a given pie chart of mutational pathways, we numerically run multiple completion experiments. Thus, we can access the distribution of the completion experiments’ results.

### A.2 Estimation experiments of unknown MPC

We evaluate two sorts of information in the framework of a laboratory experiment investigating available mutational pathways responsible for the transition from a given wild type to a given mutant phenotype. First, how many mutants do we need to observe in order to correctly estimate the total number of mutational pathways available? Secondly, some being more common than others, what does it take to correctly estimate the relative probability of each mutational pathway?

To evaluate our ability to correctly infer this information on available mutational pathways based on experimental evolution in the laboratory, we use computer simulations as numerical experiments. Indeed, that evaluation is impossible in practice since we cannot access the correct data we are trying to infer. Computer simulations allow us to start by positing what the truth could be in order to test to what extent experiments are able to recover it. For instance, we assume that four mutational pathways are available, all equally common first for simplicity (25%, 25%, 25%, 25%), and simulate the discovery of one, two, etc. Mutants, each with a single mutation corresponding to one of the pathways.

To model the fact that the true MPC is unknown, we design numerical experiments with three compartments. The MPC that experimentalists want to estimate is coded in a truth compartment. It determines the observation compartment, where randomly sampled mutants appear. We code algorithmic estimations in the modelling space based on the observed mutants. Thus, experimental estimations have no direct access to the truth compartment.

### A.3 Repeatability threshold

Consider a focal, minor mutational pathway *A* and a major mutational pathway *B,* with a target size *r* times larger. The only way for A to have a higher effective mutation rate is to have a mutational hotspot allowing it to overcome the determinism of the MPC-setup of target sizes. Thus, a higher coefficient of variation of the DMR increases stochasticity, favours mutational hotspots and decreases repeatability. In the log-normal framework for the DMR, the coefficient of variation is directly tied to parameter *σ*. Therefore, parameter *σ* of the log-normal DMR determines a threshold value *r_thresh_* above which the determinism introduced by target size is robust with 95% certainty against the stochastic aspect introduced by the DMR. Realistic values for *σ* keep this threshold below 15 (Figure 6).

## B Experimental appendix

### B.1 Description of the wrinkly spreader model system

The wrinkly spreader (WS) system in *Pseudomonas fluorescens* SBW25 is one of the best characterized experimental evolution systems and has been used to address many fundamental questions in evolutionary biology (Rainey and Travisano, 1998; Kassen et al., 2000; Beaumont et al., 2009; McDonald et al., 2009; Hammerschmidt et al., 2014; Lind et al., 2015; Gallie et al., 2019; Lind et al., 2019). The experimental setup is simple: bacteria are inoculated into static growth tubes, and competition for access to oxygen leads to strong selection for mutants with increased ability to colonize the air-liquid interface. Several types of mutants are distinguishable by their colony morphology on agar plates, which allows for isolation of many independent mutants even when present at low frequencies in a population.

The most successful phenotype is the WS morphotype, whose increased production of exopolysaccharides increases cell-cell and cell-wall adhesion (Spiers et al., 2003; Lind et al., 2016). The WS phenotype is caused by mutational activation of diguanylate-cyclases that increase the production of the second messenger c-di-GMP, a signal for biofilm formation and increased exopolysaccharide production (Goymer et al., 2006; McDonald et al., 2009; Lind et al., 2015). The WS morphotype has been found to be caused by single mutations in 19 genes in 13 regulatory pathways, all with a similar fitness increase when competed against the ancestor (Lind et al., 2015, 2019). However, mutations in some genes are rarely observed experimentally due to the small number of possible WS mutations (Lind et al., 2015). Therefore, the spectrum of WS mutations is almost completely dominated by mutations in four main pathways, *wspABCDEFR*, *awsXRO*, *mwsR* and *dgcH*. All are under negative regulation, which can be disrupted by a large number/range of mutations to produce the WS phenotype (McDonald et al., 2009; Lind et al., 2019).

It is also possible to isolate WS mutants without selecting for the ability to colonize the air-liquid interface, by using a reporter construct where mutation rates can be measured in a Luria–Delbrück fluctuation assay (McDonald et al., 2011; Lind et al., 2019). This has led to the realization that mutational hotspots play a major part in determining the outcome of experimental evolution in this model system (McDonald et al., 2009; Lind et al., 2019). *Pseudomonas protegens* Pf-5 also evolves similar morphotypes with mutations in the same main pathways (Pentz and Lind, 2021). However, it appears that mutation hotspots are not conserved between species, which could limit our ability to forecast evolution in one species based on an understanding of the GPM of another (Pentz and Lind, 2021).

### B.2 Estimation of the number of WS mutations

To examine the consequences of having major differences in the number of adaptive mutations per gene, we estimated the number of mutations that could produce the WS phenotype (Table 2). This GPM model is based on incomplete empirical data combined with an understanding of the molecular function of some of the proteins involved. It only serves to provide a realistic example of a model system with order-of-magnitude differences between the number of adaptive mutations per gene. It is not a statement about the true number of WS mutations per gene, which is difficult to measure experimentally, as we mention in the main discussion.

For the six genes with the smallest number (<5) of estimated mutations (Table 2, bottom), we base our estimates on experimentally observed mutations in a strain where the common *wsp*, *aws* and *mws* pathways have been deleted (Lind et al., 2015). Given that some mutations in these genes were observed only once, this is likely to be an underestimate.

For *dgcH,* 24 different mutations were observed (Lind et al., 2015). The mutational pattern makes it clear that there is a roughly 500 bp region where any disabling mutation, including small in-frame deletions, lead to a WS phenotype. Since many of these mutations were observed only once, we estimate that the number of possible WS mutations is 50 for *dgcH.*

Identifying all possible WS mutations in the common pathways would require characterization of thousands of independent mutants. Lacking such data, we base our estimates on the number of mutations observed (McDonald et al., 2009, 2011; Lind et al., 2019) combined with knowledge that any loss-of-function mutation in *wspF* or *awsX* produce a WS type (Bantinaki et al., 2007; McDonald et al., 2009). The *wspF* gene is about 1000 bp, so there are 3000 different possible base pair substitutions as every base can mutate to the three other. Roughly one third of these are synonymous mutations that are unlikely to have major effects on protein function, leaving 2000 base pair substitutions producing missense or nonsense mutations. Based on site-directed mutagenesis studies (Adkar et al., 2012; Jacquier et al., 2013; Firnberg et al., 2014; Lind et al., 2016; Dandage et al., 2018), we suggest that at least 5% of these 2000 mutations would severely compromise protein function, so that 100 different WS base pair substitution mutants would be possible in *wspF*. Additionally, there should be many possible small insertions and deletions that disrupt function. Thus, the 120 proposed possible WS mutations in *wspF* (Table 2) are likely to be an underestimate.

We chose to presumably underestimate the number of possible WS mutations in the common and rare pathways to be conservative about how low the median mutation rate might be. The total mutation rate to the WS phenotype, measured by fluctuation tests, is 10^-8^ per generation (Lind et al., 2019), which with 500 estimated total mutations in Table 2 gives an average mutation rate of 2 × 10^-11^ gen^-1^. Given that two mutation hotspots in *awsX* and *awsR* account for 42% of the total mutation rate (Lind et al., 2019), the median mutation rate is lower than the average.

### B.3 WS mutation rate data for the *aws* operon

The mutation rate to the WS phenotype has previously been measured in fluctuation tests using a reporter construct (Lind et al., 2019). We used the experimental data for the *aws* pathway in Appendix B (Table 3) to calibrate the log-normal DMR model.

## C Theoretical appendix

### C.1 Discussion of the MPC metrics

Here, we compare the metric that we use to summarize a given MPC by a single number, the unevenness index, to two other possible metrics, the Gini inequality coefficient and the sum of the two largest shares (SLS2). All metrics quantify the unbalance between the shares of the MPC and give a value between 0 (perfect balance) and 1 (maximal unbalance). All depend solely on the relative proportion of the MPC’s shares and are insensitive to absolute mutation rates: they quantify the unbalance of the whole MPC, not the probability of any single mutational element.

This article focuses on the unevenness index U, based on Shannon’s diversity index *H*’:

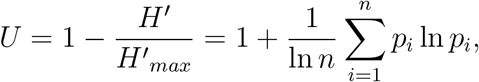

where *n* is the total number of mutational elements, and *p_i_* the probability of a particular element *i* (with all such probabilities summing to 1).

#### C.1.1 Alternative metrics

We present two alternative metrics and show that, in both cases, their range depends on the number of mutational elements, which makes them inconvenient when the total number of mutational elements may vary or be unknown as in the laboratory.

##### Gini inequality coefficient

The Gini inequality coefficient, used in economics, is based on pairwise comparisons between all mutational elements of the MPC:

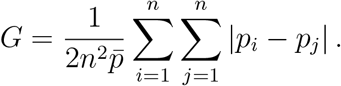

The Gini coefficient is 0 if and only if all mutation elements have the same probability. When mutations have a 100 % probability to come from a single site, it increases up to 1 – 1/*n*, which gets closer to 1 if there are many mutation sites.

##### Sum of the two largest shares

The sum of the two largest shares of an MPC is the most intuitive measure of disbalance from an experimental point of view:

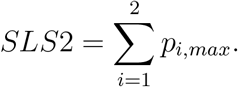

The lowest possible value for the SLS2 is 2/n, which gets closer and closer to 0 when the number of mutational elements increases. It corresponds to the MPC is perfectly balanced. The SLS2 is 1 if and only if two mutational elements take up the entire MPC.

#### C.1.2 Relationship between the different metrics

In general, each single value for a given metric corresponds to multiple possible configurations for an MPC. This variation means that there is no exact correspondence between a value of the unevenness and the other MPC metrics.

However, Figure 9 shows that this variation is moderate and that there are clear statistical correlations between the three metrics, when the number of mutations considered is fixed. All proposed MPC metrics capture in a single value the imbalance between mutation rates: the higher they are, the less equal the mutation rates. When the number of mutations increases, the Gini coefficient tends to give a more and more distorted image of the MPCs in that most MPCs with some substantial unevenness have similar, high Gini coefficients. Consequently, the Gini coefficient distinguishes balanced MPCs more effectively as well. By contrast, a high number of mutations makes the *SLS*2 correlate more linearly with the unevenness.

**Figure 9:**
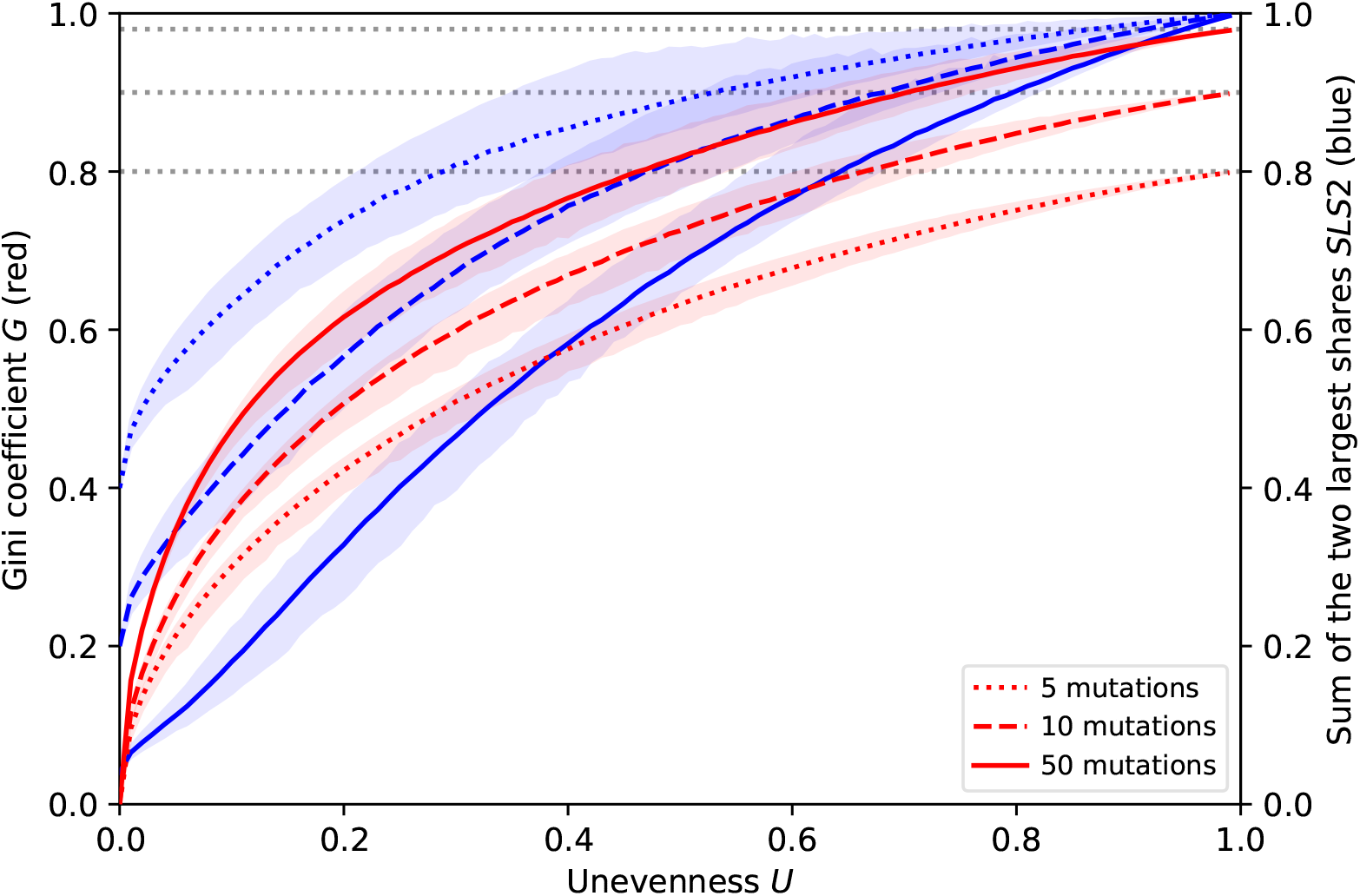
MPC metrics. Comparison of different metrics summarizing an MPC showing, for different numbers of mutations, the variation of the Gini coefficient G and of the sum of the two largest mutation rates SLS2 depending on the unevenness U, as to their mean (coloured lines), which is very close to their median, and their 95%-interval (coloured areas). The maximum Gini coefficient depends on the number of mutations (gray dotted lines).

All in all, the approach we chose in the main text could use the Gini coefficient as well. Indeed, using the distribution of any MPC metric works in principle for all. Figure 10 shows that this distribution is qualitatively similar for the unevenness and the Gini coefficient.

**Figure 10:**
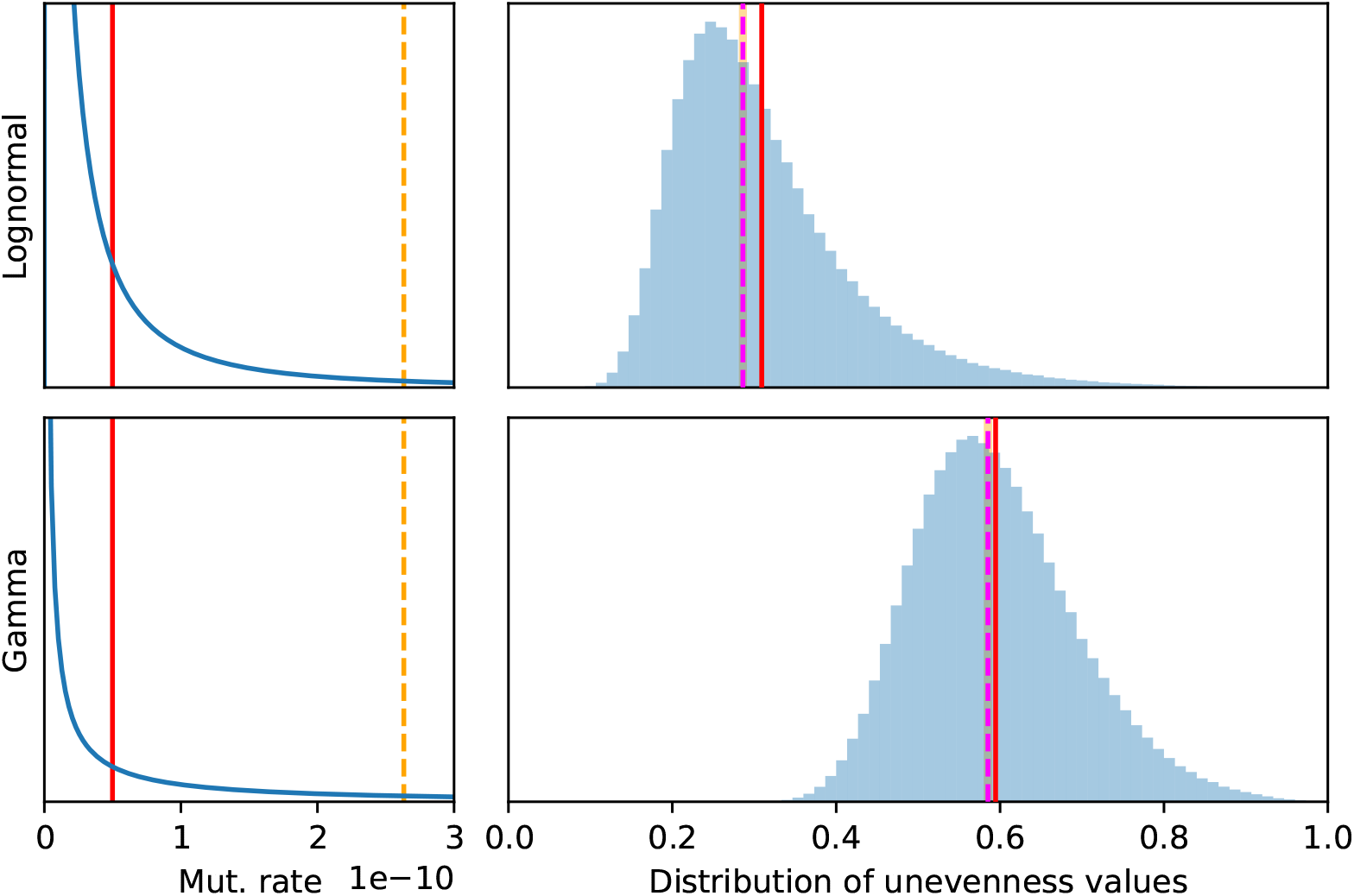
Log-normal DMR versus gamma DMR. Effect of the mathematical shape of the DMR (blue curves) on the distribution of the unevenness values (blue histograms) of 12-MPCs sampling them following the aws data setup. The DMRs have the same mean (solid red line) and standard deviation (dashed orange line), which was based on the aws calibrated log-normal DMR.

#### C.1.3 Unevenness and completion experiments’ results

##### Reproducibility with a different number of mutations

Figure 11 shows that the qualitative impact of the unevenness on the results of completion experiments is the same as in the main text even when the number of mutations considered changes.

**Figure 11:**
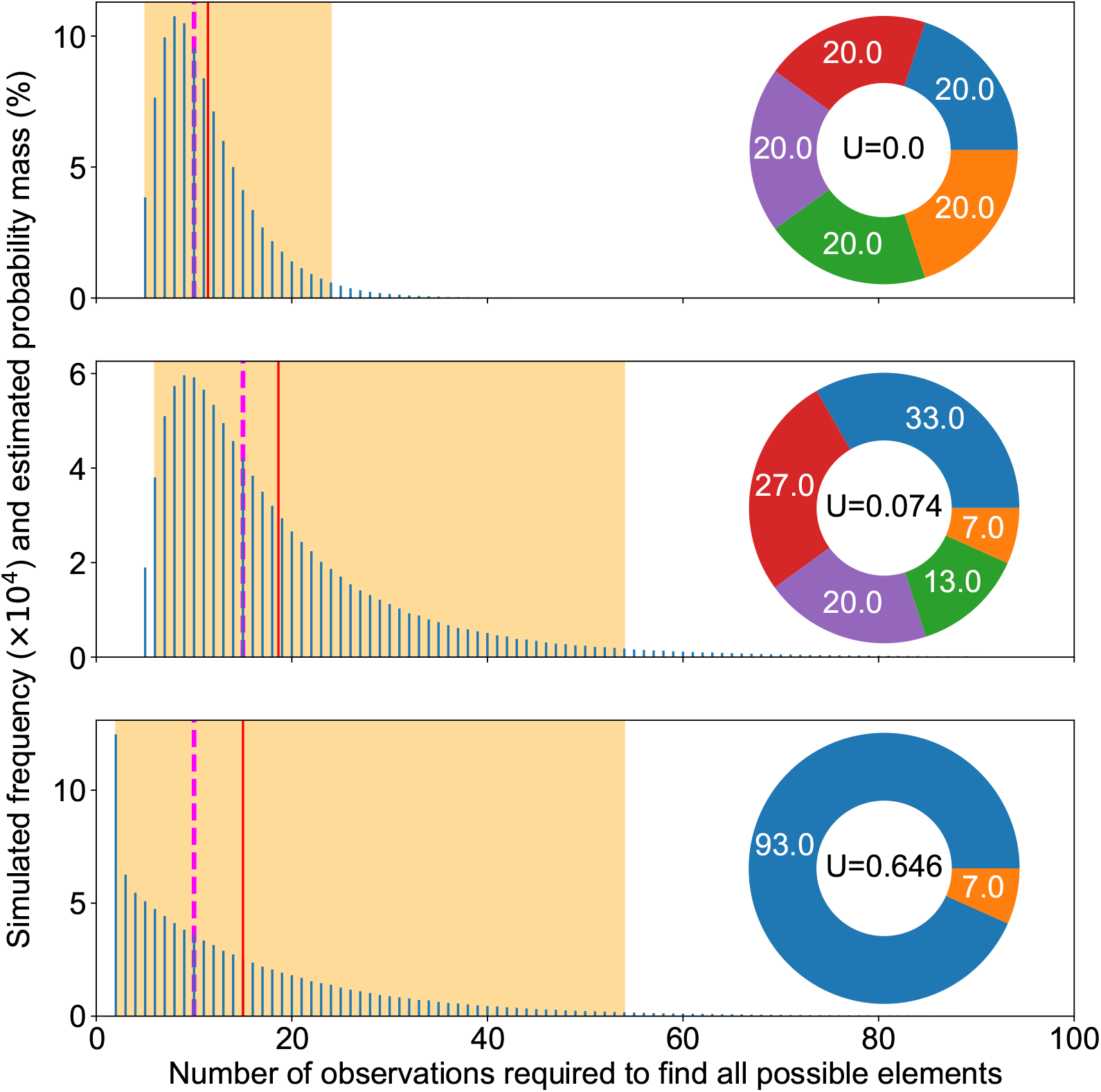
Completion experiments. Summary of completion experiments run on three examples of mutation pie charts (MPC) with different mutation rates. The smallest number of replicates needed to find all mutational elements follows a skewed distribution (blue bars), with the highest simulated value being respectively 67, 301 and 2012 out of 10^6^ experiments for each MPC. The average simulated value is shown as a solid red line, the median as a dashed magenta line and 95% of the experiment fell within the orange area.

##### The unevenness is relevant to analyze the results of completion experiments

The unevenness captures only a part of the variability of MPCs: for the same unevenness, many MPC configurations may be possible. Thus, Figure 12 shows that the variability that is not captured by the unevenness metric has a moderate impact on the variability of completion experiments. Importantly though, the median number of observations required for completion remains similar. The mean may vary more. In all cases, measures of dispersion seem to be of the same order of magnitude. By contrast, Figure 13 shows that variations in the unevenness captures dramatic changes in completion experiments’ results, which can be shifted by an order of magnitude. This not only applies to their dispersion but also to their mean and median.

**Figure 12:**
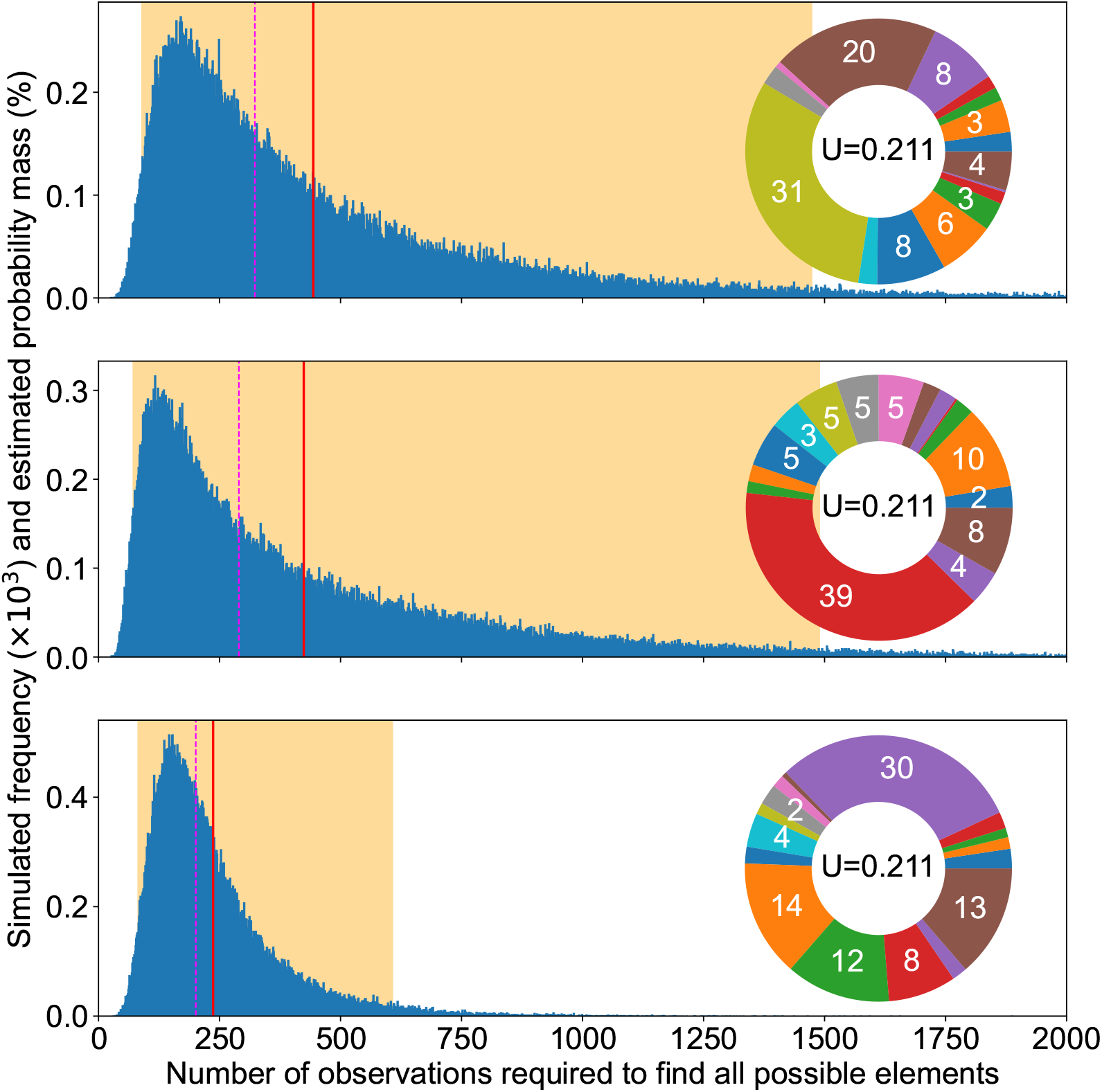
Completion experiments, variability under same unevenness. Summary of completion experiments run on three examples of 16-MPCs with different mutation rates but the same unevenness. The smallest number of replicates needed to find all mutational elements follows a skewed distribution (blue bars), with the highest simulated value being respectively 4940, 5196 and 1886 out of 10^6^ experiments for each MPC, which had Gini coefficient 0.568, 0.529 and 0.569; SS1LC 0.018, 0.044 and 0.038 (proof that the rarest is not key); SS2LC 0.010, 0.017 and 0.017; SS2MC 0.514, 0.496 and 0.445 respectively (the latter seems to correlate with the order of medians).

**Figure 13:**
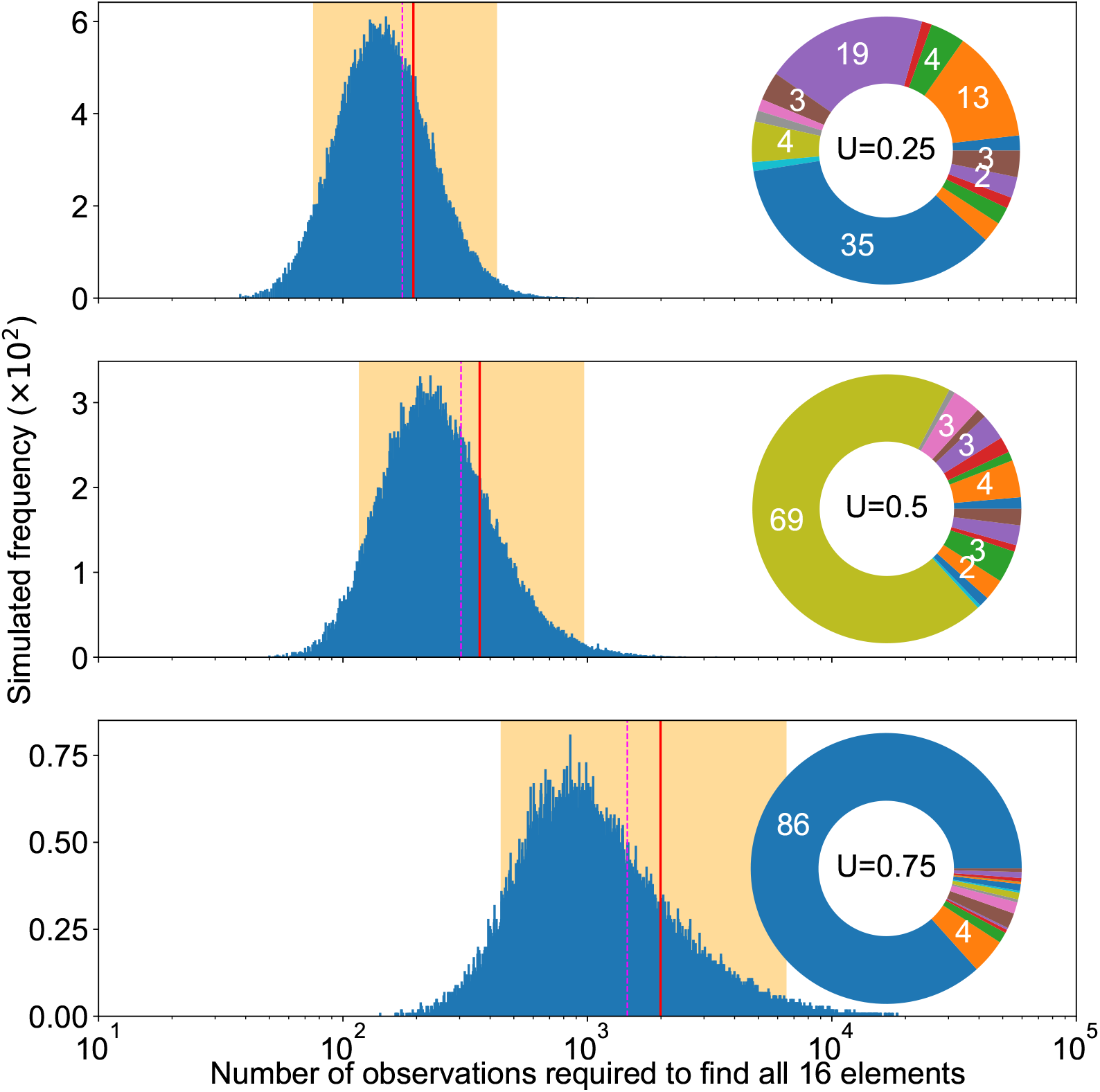
Completion experiments, variable unevenness. Summary of completion experiments run on three examples of 16-MPCs with different mutation rates and increasing unevenness. The smallest number of replicates needed to find all mutational elements follows a skewed distribution (blue bars), with the highest simulated value being respectively 1057, 3358 and 18 535 out of 10^6^ experiments for each MPC.

### C.2 Theoretical results about the particular case of an even MPC

#### C.2.1 Minimum for the expected result of completion experiments

We model the expected number E{*C_m_*} of observations to gather in order to find all possible mutations as a the expected waiting time in a generalized coupon collector’s problem. Flajolet et al. (1992) proves that:

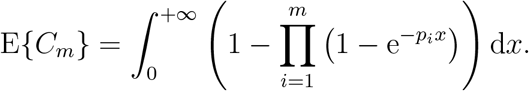

Here, we prove that, for any given number *m* ≥ 2 of mutations (shares on the pie chart), the expected value is minimal in the equiprobable case, where all mutations have the same mutation rate.

Obviously, what we want to prove is implied by the fact that, for any set 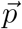 of *m* probabilities summing to 1 and for any *x* ≥ 0:

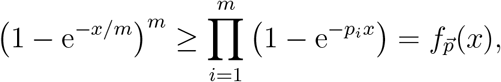

since no factor can be negative. That inequality is trivially true if any of the probabilities *p_i_* vanishes. But in general, this corresponds to a simple, convex optimization problem better formulated with a logarithm transform:

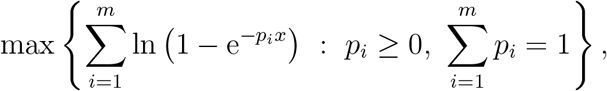

with the Lagrangian function:

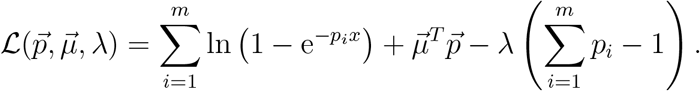

The following, necessary Karush–Kuhn–Tucker conditions provide the solution:

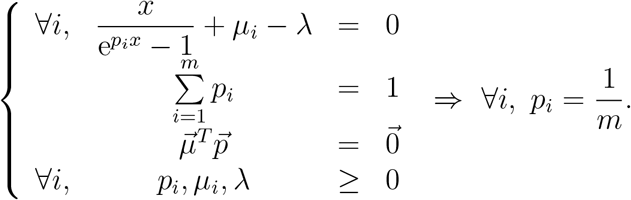

Thus, the equiprobable case minimizes the expected number *E*{*C_m_*} of observations required to find all possible mutations. Actually, an optimization approach similarly proves that the same holds for any number *j* (between 2 and m) of mutational elements to be found: the expected number of observations required is minimal in the equiprobable case.

#### C.2.2 Formulas for the case of an even MPC

In the case of an MPC with unevenness *U* = 0, the expected number of observations required to discover all mutational elements is

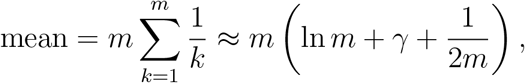

where *γ* ≈ 0.577 is the Euler-Mascheroni constant.

We can further approximate the standard deviation using the formula

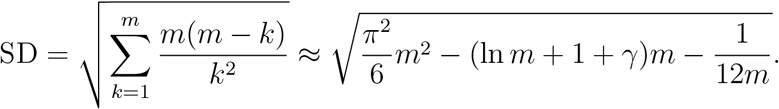

Moreover, the number of additional observations required to observe a new, *k* + 1-th element after discovering *k* different mutational elements is geometrically distributed, with expectation *m*/(*m* – *k*).

Modelling the discovery of mutational elements from an even MPC as the coupon collector’s problem in mathematics, the even case further allows for an estimation of the total number of mutational elements *m* given the number of elements already discovered *k* and the total sample size *obs* of observations gathered. Indeed, Dawkins (1991) shows that the total number *m* of mutational elements achieving the maximum likelihood satisfies

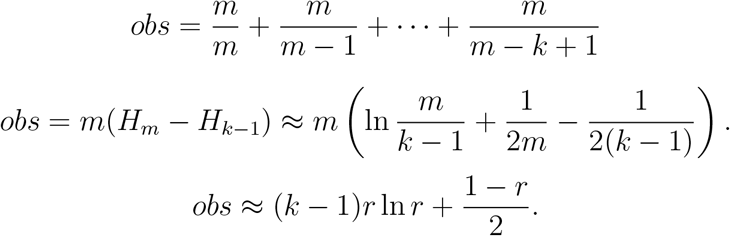

### C.3 Impact of the number of mutations on distribution of the total resulting mutation rate (TMR)

Table 4 summarizes the theoretical impact that the number of mutations has on the distribution of the total resulting mutation rate (TMR).

### C.4 Theoretical discussion on evolutionary repeatability and log-normal DMR

Our results agree with several theoretical expectations regarding the repeatability of evolution (Stoltzfus, 2021, 8.12, p. 159-161). In particular, the formalism of a log-normal DMR perfectly fits the expectation that the probability of observing parallel evolution strongly depends (main text, Figure 7) on the coefficient of variation of the DMR, modelled by parameter *σ*, and not on the absolute mutation rates, modelled by parameter μ. In the log-normal DMR framework, the probability *Pr_para_* of parallel evolution (Stoltzfus, 2021, eq. 8.8, p. 160) reduces to the simple expression:

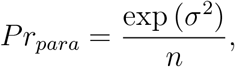

where *n* is the number of evolutionary possibilities. It allows us to interpret the aggregation of molecularly defined mutations, expected to increase evolutionary parallelism (Stoltzfus, 2021, p. 161), in terms of different scales: thus, it becomes obvious that a scale with a lower number n of mutational elements (gene or metabolic pathway, as opposed to the nucleotide level) increases the predictability of evolution, as observed experimentally (Tenaillon et al., 2012).

Our numerical quantification of the repeatability of experimental evolution illustrated the balance between the determinism of the MPC-setup and the stochasticity of the DMR-variance. It explains why experiments always observe common mutated genes, while rare pathways may only be found in an unreliable fashion among replicates, or across species. As a consequence, we should interpret mutational pathways found in several similar experiments as corresponding to a large target size (common mutations), while pathways found by some studies only should have a significantly lower target size (rare mutations). Repeatedly finding rare mutations in some studies only is in turn strong evidence that the observation span is too narrow to conclude, suggesting that all studies taken together remain underpowered.

## Notes

https://github.com/anthony-sun/Mutationrates.git

### Competing Interest Statement

The authors have declared no competing interest.

## References

B. V. Adkar, A. Tripathi, A. Sahoo, K. Bajaj, D. Goswami, P. Chakrabarti, M. K. Swarnkar, R. S. Gokhale, and R. Varadarajan. Protein model discrimination using mutational sensitivity derived from deep sequencing. Structure, 20(2):371–381, 2012. doi: 10.1016/j.str.2011.11.021.

P. Anderson and J. Roth. Spontaneous tandem genetic duplications in salmonella typhimurium arise by unequal recombination between rRNA (rrn) cistrons. Proceedings of the National Academy of Sciences, 78(5): 3113–3117, 1981. doi: 10.1073/pnas.78.5.3113.

E. Bantinaki, R. Kassen, C. G. Knight, Z. Robinson, A. J. Spiers, and P. B. Rainey. Adaptive divergence in experimental populations of pseudomonas fluorescens. III. mutational origins of wrinkly spreader diversity. Genetics, 176(1):441–453, 2007. doi: 10.1534/genetics.106.069906.

H. J. E. Beaumont, J. Gallie, C. Kost, G. C. Ferguson, and P. B. Rainey. Experimental evolution of bet hedging. Nature, 462(7269):90–93, 2009. doi: 10.1038/nature08504.

A. Boneh and M. Hofri. The coupon-collector problem revisited. Department of Computer Science Technical Reports, Purdue University, Paper 807, 1989. URL https://docs.lib.purdue.edu/cstech/807.

A. V. Cano, A. Couce, J. Masel, J. L. Payne, A. Stoltzfus, and J. F. Storz. Misrepresenting biases in arrival: a comment on svensson (2022). Eco-EvoRxiv, 2022a. doi: 10.32942/x2sg6v.

A. V. Cano, B. L. Gitschlag, H. Rozhonova, A. Stoltzfus, D. M. McCandlish, and J. L. Payne. Mutation bias and the predictability of evolution. EcoEvoRxiv, 2022b. doi: 10.32942/x2qg67.

A. V. Cano, H. Rozhoňová, A. Stoltzfus, D. M. McCandlish, and J. L. Payne. Mutation bias shapes the spectrum of adaptive substitutions. Proceedings of the National Academy of Sciences, 119(7), 2022c. doi: 10.1073/pnas.2119720119.

R. Dandage, R. Pandey, G. Jayaraj, M. Rai, D. Berger, and K. Chakraborty. Differential strengths of molecular determinants guide environment specific mutational fates. PLOS Genetics, 14(5):e1007419, 2018. doi: 10.1371/journal.pgen.1007419.

B. Dawkins. Siobhan’s problem: The coupon collector revisited. The American Statistician, 45(1):76–82, 02 1991. doi: 10.2307/2685247.

J. R. Dettman, J. L. Sztepanacz, and R. Kassen. The properties of spontaneous mutations in the opportunistic pathogen pseudomonas aeruginosa. BMC Genomics, 17(1), 2016. doi: 10.1186/s12864-015-2244-3.

G. C. Ferguson, F. Bertels, and P. B. Rainey. Adaptive divergence in experimental populations of pseudomonas fluorescens. v. insight into the niche specialist fuzzy spreader compels revision of the model pseudomonas radiation. Genetics, 195(4):1319–1335, 2013. doi: 10.1534/genetics.113.154948.

M. Ferrante and M. Saltalamacchia. The coupon collector’s problem Materials Matemàtics, 2014(2), 2014. ISSN 1887-1097. URL https://mat.uab.cat/web/matmat/wp-content/uploads/sites/23/2020/05/v2014n02.pdf.

E. Firnberg, J. W. Labonte, J. J. Gray, and M. Ostermeier. A comprehensive, high-resolution map of a gene’s fitness landscape. Molecular Biology and Evolution, 31(6):1581–1592, 2014. doi: 10.1093/molbev/msu081.

P. Flajolet, D. Gardy, and L. Thimonier. Birthday paradox, coupon collectors, caching algorithms and self-organizing search. Discrete Applied Mathematics, 39(3):207–229, 1992. ISSN 0166-218X. doi: 10.1016/0166-218X(92)90177-C.

P. L. Foster. Methods for determining spontaneous mutation rates. In J. L. Campbell and P. Modrich, editors, DNA Repair, Part B, volume 409 of Methods in Enzymology, pages 195–213. Elsevier, 2006. doi: 10.1016/s0076-6879(05)09012-9.

P. L. Foster, H. Lee, E. Popodi, J. P. Townes, and H. Tang. Determinants of spontaneous mutation in the bacterium escherichia coli as revealed by whole-genome sequencing. Proceedings of the National Academy of Sciences, 112(44), 2015. doi: 10.1073/pnas.1512136112.

E. Furman, D. Hackmann, and A. Kuznetsov. On log-normal convolutions: An analytical-numerical method with applications to economic capital determination. Insurance: Mathematics and Economics, 90:120–134, 2020. doi: 10.1016/j.insmatheco.2019.10.003.

J. Gallie, F. Bertels, P. Remigi, G. C. Ferguson, S. Nestmann, and P. B. Rainey. Repeated phenotypic evolution by different genetic routes in pseudomonas fluorescens SBW25. Molecular Biology and Evolution, 36(5):1071–1085, 2019. doi: 10.1093/molbev/msz040.

K. Gomez, J. Bertram, and J. Masel. Mutation bias can shape adaptation in large asexual populations experiencing clonal interference. Proceedings of the Royal Society B: Biological Sciences, 287(1937):20201503, 2020. doi: 10.1098/rspb.2020.1503.

P. Goymer, S. G. Kahn, J. G. Malone, S. M. Gehrig, A. J. Spiers, and P. B. Rainey. Adaptive divergence in experimental populations of pseudomonas fluorescens. II. role of the GGDEF regulator WspR in evolution and development of the wrinkly spreader phenotype. Genetics, 173(2):515–526, 2006. doi: 10.1534/genetics.106.055863.

D. L. Halligan and P. D. Keightley. Spontaneous mutation accumulation studies in evolutionary genetics. Annual Review of Ecology, Evolution, and Systematics, 40(1):151–172, 2009. doi: 10.1146/annurev.ecolsys.39.110707.173437.

K. Hammerschmidt, C. J. Rose, B. Kerr, and P. B. Rainey. Life cycles, fitness decoupling and the evolution of multicellularity. Nature, 515(7525):75–79, 2014. doi: 10.1038/nature13884.

J. S. Horton, L. M. Flanagan, R. W. Jackson, N. K. Priest, and T. B. Taylor. A mutational hotspot that determines highly repeatable evolution can be built and broken by silent genetic changes. Nature Communications, 12 (1), 2021. doi: 10.1038/s41467-021-26286-9.

H. Jacquier, A. Birgy, H. Le Nagard, Y. Mechulam, E. Schmitt, J. Glodt, B. Bercot, E. Petit, J. Poulain, G. Barnaud, P.-A. Gros, and O. Tenaillon. Capturing the mutational landscape of the beta-lactamase TEM-1. Proceedings of the National Academy of Sciences, 110(32):13067–13072, 2013. doi: 10.1073/pnas.1215206110.

R. Kassen, A. Buckling, G. Bell, and P. B. Rainey. Diversity peaks at intermediate productivity in a laboratory microcosm. Nature, 406(6795): 508–512, 2000. doi: 10.1038/35020060.

A. Knöppel, M. Knopp, L. M. Albrecht, E. Lundin, U. Lustig, J. Näsvall, and D. I. Andersson. Genetic adaptation to growth under laboratory conditions in escherichia coli and salmonella enterica. Frontiers in Microbiology, 9, 2018. doi: 10.3389/fmicb.2018.00756.

S. Koskiniemi, S. Sun, O. G. Berg, and D. I. Andersson. Selection-driven gene loss in bacteria. PLoS Genetics, 8(6):e1002787, 2012. doi: 10.1371/journal.pgen.1002787.

P. A. Lind and D. I. Andersson. Whole-genome mutational biases in bacteria. Proceedings of the National Academy of Sciences, 105(46):17878–17883, 2008. doi: 10.1073/pnas.0804445105.

P. A. Lind, A. D. Farr, and P. B. Rainey. Experimental evolution reveals hidden diversity in evolutionary pathways. eLife, 4:e07074, 2015. ISSN 2050-084X. doi: 10.7554/eLife.07074.

P. A. Lind, A. D. Farr, and P. B. Rainey. Evolutionary convergence in experimental pseudomonas populations. The ISME Journal, 11(3):589–600, 2016. doi: 10.1038/ismej.2016.157.

P. A. Lind, E. Libby, J. Herzog, and P. B. Rainey. Predicting mutational routes to new adaptive phenotypes. eLife, 8:e38822, 2019. ISSN 2050-084X. doi: 10.7554/eLife.38822.

H. Long, S. Kucukyildirim, W. Sung, E. Williams, H. Lee, M. S. Ackerman, T. G. Doak, H. Tang, and M. Lynch. Background mutational features of the radiation-resistant bacterium deinococcus radiodurans. Molecular Biology and Evolution, 32(9):2383–2392, 2015. doi: 10.1093/molbev/msv119.

H. Long, W. Sung, S. Kucukyildirim, E. Williams, S. F. Miller, W. Guo, C. Patterson, C. Gregory, C. Strauss, C. Stone, C. Berne, D. Kysela, W. R. Shoemaker, M. E. Muscarella, H. Luo, J. T. Lennon, Y. V. Brun, and M. Lynch. Evolutionary determinants of genome-wide nucleotide composition. Nature Ecology and Evolution, 2(2):237–240, 2018. doi: 10.1038/s41559-017-0425-y.

S. E. Luria and M. Delbrück. Mutations of bacteria from virus sensitivity to virus resistance. Genetics, 28(6):491–511, 1943. doi: 10.1093/genetics/28.6.491.

R. C. MacLean, G. G. Perron, and A. Gardner. Diminishing returns from beneficial mutations and pervasive epistasis shape the fitness landscape for rifampicin resistance in pseudomonas aeruginosa. Genetics, 186(4): 1345–1354, 2010. doi: 10.1534/genetics.110.123083.

D. M. McCandlish and A. Stoltzfus. Modeling evolution using the probability of fixation: History and implications. The Quarterly Review of Biology, 89 (3):225–252, 2014. doi: 10.1086/677571.

M. J. McDonald, S. M. Gehrig, P. L. Meintjes, X.-X. Zhang, and P. B. Rainey. Adaptive divergence in experimental populations of pseudomonas fluorescens. IV. genetic constraints guide evolutionary trajectories in a parallel adaptive radiation. Genetics, 183(3):1041–1053, 2009. doi: 10.1534/genetics.109.107110.

M. J. McDonald, T. F. Cooper, H. J. E. Beaumont, and P. B. Rainey. The distribution of fitness effects of new beneficial mutations in pseudomonas fluorescens. Biology Letters, 7(1):98–100, 2011. doi: 10.1098/rsbl.2010.0547.

J. G. Monroe, T. Srikant, P. Carbonell-Bejerano, C. Becker, M. Lensink, M. Exposito-Alonso, M. Klein, J. Hildebrandt, M. Neumann, D. Kliebenstein, M.-L. Weng, E. Imbert, J. Ågren, M. T. Rutter, C. B. Fenster, and D. Weigel. Mutation bias reflects natural selection in arabidopsis thaliana. Nature, 602(7895):101–105, 2022. doi: 10.1038/s41586-021-04269-6.

A. W. Murray. Can gene-inactivating mutations lead to evolutionary novelty? Current Biology, 30(10):R465–R471, 2020. doi: 10.1016/j.cub.2020.03.072.

H. B. Nath. On the collector’s sequential sample size. Trabajos de estadistica y de investigatión operativa, 25(3):85–88, 1974. doi: 10.1007/bf03005894.

J. Pan, W. Li, J. Ni, K. Wu, I. Konigsberg, C. E. Rivera, C. Tincher, C. Gregory, X. Zhou, T. G. Doak, H. Lee, Y. Wang, X. Gao, M. Lynch, and H. Long. Rates of mutations and transcript errors in the foodborne pathogen salmonella enterica subsp. enterica. Molecular Biology and Evolution, 39(4), 2022. doi: 10.1093/molbev/msac081.

T. J. Pentz and P. A. Lind. Forecasting of phenotypic and genetic outcomes of experimental evolution in pseudomonas protegens. PLOS Genetics, 17 (8):1–24, 08 2021. doi: 10.1371/journal.pgen.1009722.

P. B. Rainey and M. Travisano. Adaptive radiation in a heterogeneous environment. Nature, 394(6688):69–72, 1998. doi: 10.1038/27900.

T. S. Sankar, B. D. Wastuwidyaningtyas, Y. Dong, S. A. Lewis, and J. D. Wang. The nature of mutations induced by replication-transcription collisions. Nature, 535(7610):178–181, 2016. doi: 10.1038/nature18316.

J. W. Schroeder, P. Yeesin, L. A. Simmons, and J. D. Wang. Sources of spontaneous mutagenesis in bacteria. Critical Reviews in Biochemistry and Molecular Biology, 53(1):29–48, 2017. doi: 10.1080/10409238.2017.1394262.

A. J. Spiers, J. Bohannon, S. M. Gehrig, and P. B. Rainey. Biofilm formation at the air-liquid interface by the pseudomonas fluorescens SBW25 wrinkly spreader requires an acetylated form of cellulose. Molecular Microbiology, 50(1):15–27, 2003. doi: 10.1046/j.1365-2958.2003.03670.x.

A. Stoltzfus. Mutation, Randomness, and Evolution. Oxford University Press, 2021. ISBN 978-0-19-884445-7. doi: 10.1093/oso/9780198844457.001.0001.

A. Stoltzfus and D. M. McCandlish. Mutational biases influence parallel adaptation. Molecular Biology and Evolution, 34(9):2163–2172, 2017. doi: 10.1093/molbev/msx180.

W. Sung, M. S. Ackerman, J.-F. Gout, S. F. Miller, E. Williams, P. L. Foster, and M. Lynch. Asymmetric context-dependent mutation patterns revealed through mutation-accumulation experiments. Molecular Biology and Evolution, 32(7):1672–1683, 2015. doi: 10.1093/molbev/msv055.

W. Sung, M. S. Ackerman, M. M. Dillon, T. G. Platt, C. Fuqua, V. S. Cooper, and M. Lynch. Evolution of the insertion-deletion mutation rate across the tree of life. G3 Genes |Genomes| Genetics, 6(8):2583–2591, 2016. doi: 10.1534/g3.116.030890.

E. I. Svensson. Multivariate selection and the making and breaking of mutational pleiotropy. Evolutionary Ecology, 36(5):807–828, 2022. doi: 10.1007/s10682-022-10195-4.

E. I. Svensson and D. Berger. The role of mutation bias in adaptive evolution. Trends in Ecology and Evolution, 34(5):422–434, 2019. doi: 10.1016/j.tree.2019.01.015.

O. Tenaillon, A. Rodríguez-Verdugo, R. L. Gaut, P. McDonald, A. F. Bennett, A. D. Long, and B. S. Gaut. The molecular diversity of adaptive convergence. Science, 335(6067):457–461, 2012. doi: 10.1126/science.1212986.

D. van Ditmarsch, K. E. Boyle, H. Sakhtah, J. E. Oyler, C. D. Nadell, E. Déziel, L. E. P. Dietrich, and J. B. Xavier. Convergent evolution of hyperswarming leads to impaired biofilm formation in pathogenic bacteria. Cell Reports, 4(4):697–708, 2013. doi: 10.1016/j.celrep.2013.07.026.

L. Y. Yampolsky and A. Stoltzfus. Bias in the introduction of variation as an orienting factor in evolution. Evolution and Development, 3(2):73–83, 2001. doi: 10.1046/j.1525-142x.2001.003002073.x.

